# Methylation status of *VTRNA2-1*/*nc886* is stable across human populations, monozygotic twin pairs and in majority of somatic tissues

**DOI:** 10.1101/2022.06.21.496995

**Authors:** Saara Marttila, Hely Tamminen, Sonja Rajić, Pashupati P Mishra, Terho Lehtimäki, Olli Raitakari, Mika Kähönen, Laura Kananen, Juulia Jylhävä, Sara Hägg, Thomas Delerue, Annette Peters, Melanie Waldenberger, Marcus E Kleber, Winfried März, Riitta Luoto, Jani Raitanen, Elina Sillanpää, Eija K Laakkonen, Aino Heikkinen, Miina Ollikainen, Emma Raitoharju

## Abstract

**Aims and methods:** Our aim was to characterise the methylation level of a polymorphically imprinted gene, *VTRNA2-1*/*nc886*, in human populations and somatic tissues. We utilised 48 datasets, consisting of >30 different tissues and >30 000 individuals.

**Results:** We show that the *nc886* methylation status is associated with twin status and ethnic background, but the variation between populations is limited. Monozygotic twin pairs present concordant methylation, while ∼30% of dizygotic twin pairs present discordant methylation in the *nc886* locus. The methylation levels of *nc886* are uniform across somatic tissues, except in cerebellum and skeletal muscle.

**Conclusion:** We hypothesize that the *nc886* imprint is established in the oocyte and that after implantation, the methylation status is stable, excluding a few specific tissues.

## Introduction

Genomic imprinting can be defined as the expression of a gene only from the maternal or the paternal allele, while the corresponding allele in the other chromosome is silenced via epigenetic mechanisms, including DNA methylation (1). The epigenetic profiles maintaining the imprinted status are established during gametogenesis when first the DNA methylation pattern is erased, after which the parent-of-origin-related DNA methylation pattern is created. For males, the DNA methylation profile of the sperm, including imprints, is completed in the primordial germ cells, before the birth of the male child. In oocytes, on the other hand, *de novo* DNA methylation will begin only after the birth of the female child, during the follicular growth phase, with gene-specific timepoints for imprinting reported (2–4). Canonically imprinted genes, approximately 130 of which exist in humans, retain the parent-of-origin-related expression pattern throughout an individualś life in all of their somatic tissues, although tissue or developmental stage-specific imprinting can be seen, for example, in the placenta (5–7). The significance of intact genetic imprints is highlighted by the severe disorders caused by imprinting defects (8).

In humans, the locus harbouring non-coding 886 (*nc886*, also known as *VTRNA2-1*) is polymorphically imprinted, with approximately 75% of individuals having a methylated maternal allele, i.e. imprinted *nc886* locus, and the remaining approximately 25% having both maternal and paternal *nc886* allele unmethylated (9–11). According to current literature, this pattern is not due to genetic variation (9,12,13). We have also previously identified individuals who escape this bimodal methylation pattern. We found they present either intermediate methylation levels (methylation beta value 0.20-0.40 i.e. methylation level of 20-40%, in approximately 1-5% of the population) or methylation beta values over 0.60 (methylation level of over 60%), indicating that also the paternal allele has gained methylation in somatic tissues (in approximately 0.1% of the population) (11). The *nc886* locus codes for a 102nt long non-coding RNA, which might then be cleaved into miRNA-like short non-coding RNAs, although the nature of these RNAs is still widely debated (14,15).

The establishment of the *nc886* imprint has been suggested to be an early event in the zygote, happening between four and six days after fertilization (13). Recently, it was suggested that the methylation pattern of *nc886* is already established in the preconceptional oocyte (10). Early establishment of *nc886* methylation status is supported by the fact that it has been shown to be uniform across analysed somatic tissues (10,16,17). As the methylation pattern of *nc886* had been reported to be concordant in monozygotic twin pairs (MZ) but not in dizygotic twin pairs (DZ), genetic factors were hypothesized to influence the methylation pattern (9), which was later shown not to be the case (10,11,18). Once established, the methylation status of *nc886* is stable from childhood to adolescence (16) and from adolescence to adulthood (11).

Changes in the proportion of individuals with methylated or unmethylated maternal *nc886* allele have been associated with maternal age (9,11), maternal socioeconomic status (11) and maternal alcohol consumption (10), as well as the season of conception in rural Gambia (9,16). Furthermore, the methylation level or status of *nc886* or level of nc886 RNAs transcribed from the locus have been associated with childhood BMI (19), adiposity and cholesterol levels (11), as well as allergies (20), asthma (21), infections (22) and inflammation (23). Interestingly, the *nc886* methylation status and the level of nc886 RNAs have also been associated with indicators of glucose metabolism (11,24). These findings suggest that the methylation status of *nc886* is a potential molecular mediator of the Developmental Origins of Health and Disease (DOHaD) hypothesis (also known as the Barker hypothesis) (25).

Detailed understanding of the determinants and functions of the methylation status of *nc886* is still lacking. *In vitro* methods are of limited feasibility, as both carcinogenesis (13) and pluripotency induction (11) have an effect on the methylation pattern at the *nc886* locus.

Currently, no animal models are available to study *nc886*, as most species, including mice and rats, do not harbour the gene, and in species harbouring the *nc886* gene, the locus is not polymorphically imprinted (26). Thus, we wanted to utilise available resources, the numerous existing DNA methylation datasets from humans, to gain insight on the methylation of *nc886*, a unique polymorphically imprinted gene and a potential molecular mediator of DOHaD.

More precisely, we wanted to investigate 1) the prevalence of *nc886* DNA methylation status groups in a large number of human populations (with a total n > 30 000) with divergent historical and geographical origins in order to identify the factors associating with the existing variation, 2) the *nc886* methylation status in MZ and DZ twin pairs to elucidate the contribution of shared gametes vs unique gametes in shared prenatal environment to the establishment of *nc886* methylation pattern, and finally 3) the *nc886* methylation patterns in a larger variety of tissues, including brain regions and placenta, which have been shown to express a multitude of imprinted genes, as well as present dynamic imprinting (5,27,28) to have insights on the potential function of *nc886* and the stability of this polymorphic imprint in human tissues.

## Materials and Methods

### Datasets

In this study, we utilized 48 DNA methylation datasets, including DILGOM, FTC, ERMA, KORA, LURIC, NELLI, SATSA and YFS as well as 39 datasets available in Gene Expression Omnibus (GEO) (29) consisting of >30 different tissues and >30 000 individuals. (Supplementary Table 1). These datasets were used to study the methylation of *nc886* locus across populations, in twin pairs and across different tissues, with some datasets utilized in multiple settings.

For the population analysis, we included 32 datasets, with the number of individuals ranging from 131 to 2711 (total n = 30 347). In these datasets, the DNA methylation data was available from blood (11,30–48), separated peripheral blood cells (49–51), blood spots (19), umbilical cord buffy coats (52), foetal cord tissue (53) or buccal swabs (52). Association of zygosity (MZ vs DZ pairs) with *nc886* methylation was analysed in 5 datasets (36,47,48,54,55).

To analyse the methylation of *nc886* across different tissues, 17 datasets were utilized. These included a dataset consisting of 30 tissues from a 112-year-old female (56), as well as datasets consisting of different brain regions ((57), GSE134379), adipose tissue (54,58), muscle (46,55–57, GSE142141, GSE171140), liver (58), buccal swabs (52,61), skin (62), sperm (63,64) and placenta (65–68). All utilized datasets are described in detail in Supplementary Table 1.

DILGOM (the Dietary, Lifestyle, and Genetic Determinants of Obesity and Metabolic Syndrome) (44) has been collected as an extension of the FINRISK 2007 survey. FINRISK surveys consist of cross-sectional, population-based studies conducted to monitor the risk of chronic diseases in Finland (69). The data used for the research were obtained from the THL biobank (study number THLBB2021_22).

FTC (the Finnish Twin Cohort) study includes three longitudinal cohorts (70,71) - the Older Twin cohort (71), FinnTwin16 (FT16) (72), and FinnTwin12 (FT12) (73). In this study, we included two subsets of FT16 and FT12 cohorts - a smaller subset of individuals for whom DNA methylation data was available from muscle, adipose tissue and blood (60) and a larger subset with methylation data only available from blood (54). For 49 MZ twin pairs, the information on whether they were dichorionic and diamniotic (DCDA, 22 pairs), monochorionic and diamniotic (MCDA, 9 pairs), or monochorionic and monoamniotic (MCMD, 18 pairs) was available. Participants of FTC were born before 1987 in Finland.

ERMA (Estrogenic Regulation of Muscle Apoptosis) study was a prospective cohort study designed to reveal how hormonal differences over the menopausal stages affect the physiological and psychological functioning of middle-aged women (74). The original ERMA cohort includes 47- to 55-year-old women living in the city of Jyväskylä or neighbouring municipalities in Finland. For this study, a subset of 47 individuals with both whole blood and muscle tissue samples available were included (60).

KORA (Kooperative Gesundheitsforschung in der Region Augsburg/the Cooperative Health Research in the Region Augsburg cohort) is a population-based health survey that collected both clinical and genetic data from the general population in the region of Augsburg and two surrounding counties in Germany (75). Data utilized here is from KORA F4 (collected in 2006-2008) and KORA FF4 (collected in 2013-2014) cohorts.

LURIC (the LUdwigshafen RIsk and Cardiovascular Health study) consists of patients of German ancestry who underwent coronary angiography between 1997 and 2000 at a tertiary care centre in Southwestern Germany (76). For this study, we utilized 2423 individuals with DNA methylation data available.

NELLI (Neuvonta, ELintavat ja LIikunta neuvolassa) cohort consists of pregnant mothers participating in an intervention study aimed at preventing gestational diabetes from 14 municipalities in the southern part of Finland (77). Included in this study is data from children of these mothers, collected at a 7-year follow-up (61). The children participating in this study were born in 2007/2008 in Finland

SATSA (the Swedish Adoption/Twin Study of Aging) is a population-based study drawn from the Swedish Twin Registry that includes same-sex pairs of twins reared together and pairs separated before the age of 11, collected between 1984 and 2014 (78), 478 of whom had DNA methylation data available.

YFS (Young Finns Study) is a multicentre follow-up study on cardiovascular risk from childhood to adulthood, launched in 1980 (79). DNA methylation data utilized here is from the 30-year follow-up in 2011, including 1714 individuals. The participants in this study were born between 1962 and 1977 in Finland.

### DNA methylation data processing

The majority of datasets in GEO were available as processed data, and these datasets were utilized as such. Datasets GSE61454, GSE71678, GSE134379, and GSE157896 were downloaded as raw idat-files, extracted with minfi package function read.metharray.exp and quantile normalized with default settings (80). For DILGOM (44), FTC (54,60), ERMA (60), KORA (11,45), LURIC (46), SATSA (47) and YFS (11), the processing of DNA methylation data has been described in detail in previous publications referenced here. For all datasets, information on sample material, Illumina array type (Illumina Infinium MethylationEPIC or Methylation450K BeadChip), and the used processing methods are provided in Supplementary Table 1.

For NELLI, genomic DNA from buccal swabs was extracted with Gentra Buccal Cell Kit (QIAGEN, cat.no: 158845) and stored at -20°C. Aliquots of 1 μg of DNA were subjected to bisulphite conversion, and a 4-μl aliquot of bisulphite-converted DNA was subjected to whole-genome amplification, followed by enzymatic fragmentation and hybridization onto an Illumina Infinium MethylationEPIC BeadChip at Helmholtz Zentrum, Munich, Germany. The arrays were scanned with the iScan reader (Illumina). All the analysed samples had a sum of detection p-values across all the probes less than 0.01. Logged (log2) median of methylated and unmethylated intensities of the analysed samples clustered well visually except for one outlier which was excluded from the analysis. Samples were checked for discrepancies between the reported and the predicted sex and none failed the test. Normalization was done with a stratified quantile normalization method implemented in preprocessQuantile function in minfi R/Bioconductor package (80,81). Probes with a detection p-value of more than 0.01 in 99% of the samples were filtered out. Similarly, cross-reactive probes and probes with SNPs were excluded from the analysis (82,83). All pre-processing steps were performed using functions implemented in the minfi R/Bioconductor package (80).

### Clustering of individuals according to nc886 locus methylation level

For all datasets utilised, we retrieved the methylation values for 14 CpGs - cg07158503, cg04515200, cg13581155, cg11978884, cg11608150, cg06478886, cg04481923, cg18678645, cg06536614, cg25340688, cg26896946, cg00124993, cg08745965, and cg18797653, shown to display bimodal DNA methylation pattern in the *nc886* locus, which is indicative of polymorphic imprinting (9,11). For some datasets, the methylation data was not available from all 14 CpGs as, depending on the quality control steps performed, some probes had been omitted from the data. The number of probes available for each dataset is provided in Supplementary Table 1.

Based on the methylation levels of the 14 CpGs, individuals were clustered into 3 groups by k-means clustering. Based on the median methylation level of the cluster, *nc886* methylation status groups were defined as imprinted (typical methylation β value > 0.40, indicative of monoallelic methylation), non-methylated (typical methylation β value < 0.15, indicative of two unmethylated alleles), and intermediately methylated (typical methylation β value 0.15-0.40). Data was visualized to verify that the clustering had identified the groups as expected. The intermediately methylated individuals could not be detected in all datasets. In the datasets they were identified, the proportion ranged from 2% to 6%.

To verify that the clustering is reproducible across different datasets, we compared the clustering results within one dataset processed with different methods. We established that while the imprinted group could be reliably identified across different normalization methods, there were some inconsistencies between intermediately methylated individuals and non-methylated individuals (Supplementary data 1). Therefore, for analyses where proportions of *nc886* status groups were investigated across datasets, we combined the intermediately methylated and non-methylated clusters, i.e. *nc886* status is described as ‘imprinted’ or as ‘other’, the latter group including both intermediately methylated and non-methylated individuals. Only when comparing individuals within a dataset, namely in the case of MZ twin pairs, we have kept all three status groups in the analyses.

### Comparison to established imprinted genes

When analysing different tissues, we also examined the methylation level of six known imprinted genes (*DIRAS3*, *KCNQ10T1*, *MEG3*, *MEST*, *NAP1L5*, *PEG10*, and *ZNF597*) (84). This was done in tissues not presenting bimodal *nc886* pattern, to ensure imprinted genes, in general, do not display atypical methylation patterns in these tissues.

### Statistical analyses

Differences in the proportion of individuals with imprinted *nc886* locus between sexes or in a case-control setting (depression (GSE125105), assisted reproductive technologies (GSE157896), rheumatoid arthritis (GSE42861), gestational diabetes mellitus (GSE141065), schizophrenia (GSE80417, GSE84727), inflammatory bowel disease (GSE87648), childhood abuse (GSE132203), and Parkinson’s disease (GSE111629)) were investigated with χ^2^-test, with a threshold for significance set at p < 0.05 (Supplementary Table 2). For DZ twin pairs, the mathematically estimated proportion of miss-matched pairs (i.e. one twin imprinted, one intermediately methylated or non-methylated) was calculated. The difference between the observed number of miss-matched pairs and the estimated numbers was then investigated with the χ^2^-test, with a threshold for significance set at p < 0.05.

In tissues not presenting bimodal *nc886* methylation pattern, the difference in methylation levels between imprinted and other groups (as clustered according to a tissue presenting the expected methylation pattern from the same individuals) were analysed, with Mann-Whitney U-test with the threshold for significance set at p < 0.05. The median methylation levels in the *nc886* locus were correlated between different tissues with Spearman correlation, with the threshold for significance set at p <0.05.

## Results

### Methylation status of nc886 across population cohorts

We characterized the methylation status of the *nc886* locus in 32 cohorts, including both population-based and case-control settings, consisting of 30 347 individuals in total. In the majority of the cohorts, the participants were described as being of European descent, white or Caucasian, hereafter referred to as white. In the datasets utilized, DNA methylation data was from whole blood, blood cells or blood cell subtypes, buccal swabs or foetal cord tissue (Supplementary Table 2). In these tissues, the methylation level of the *nc886* locus followed the expected bimodal pattern, and thus the individuals could be clustered into *nc886* methylation status groups (Supplementary Figure 1). Across all datasets, the proportion of imprinted individuals (individuals with the methylation level indicative of monoallelic methylation) varied between 65.8% and 83.5%, with an average percentage of 75.3% (Figure 1, Supplementary Table 2). When considering only cohorts consisting of white singletons, the proportion of imprinted individuals varied less, and was between 72.6% and 77.6% (Figure 1, Supplementary Table 2).

**Figure 1.**
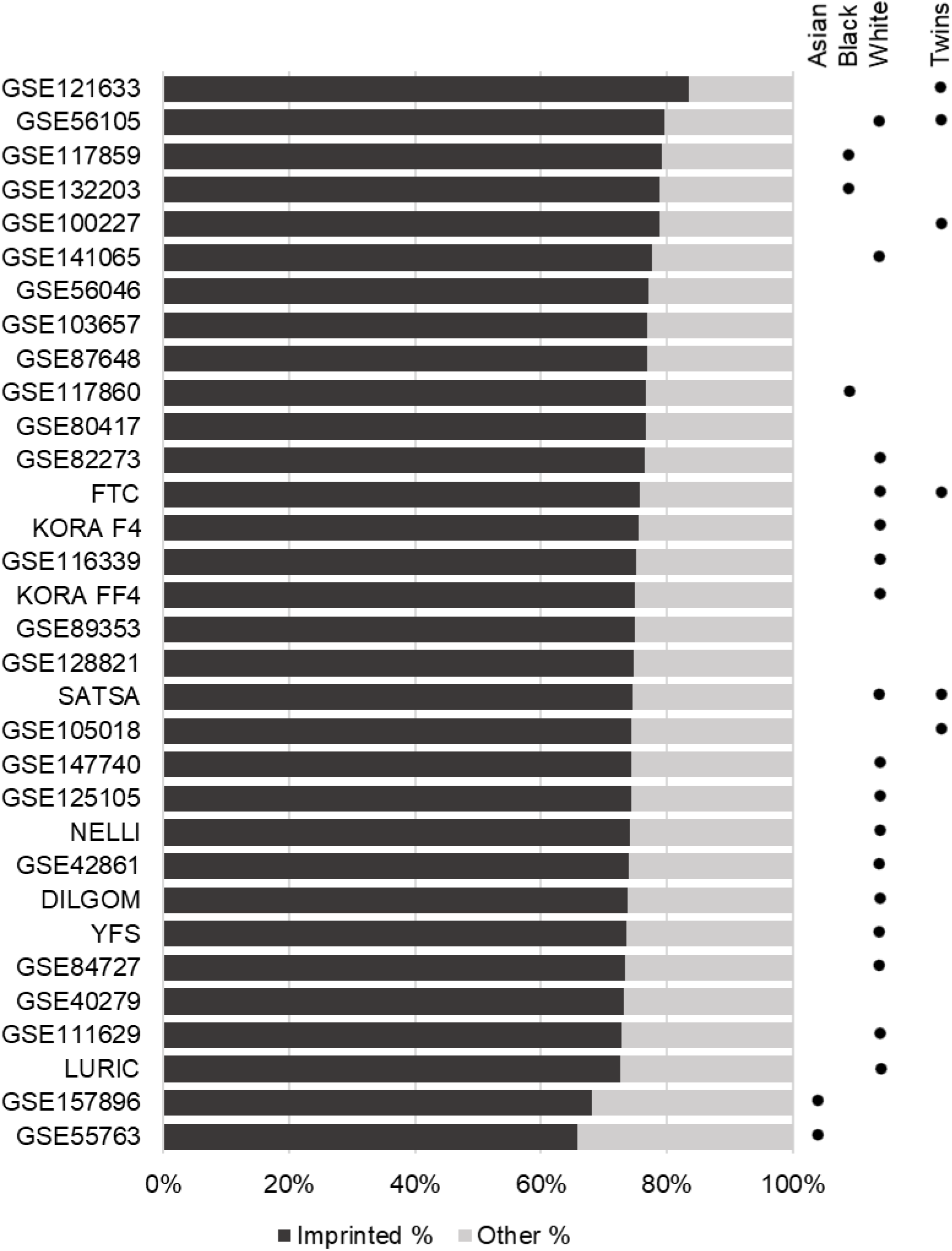
Proportion of imprinted individuals across cohorts utilized in this study. Individuals were clustered as imprinted and other (including non-methylated and intermediately methylated). Ethnicity was not specified for all cohorts utilized. Details of each cohort can be found from Supplementary Tables 1 and 2.

The lowest proportion of imprinted individuals was observed in datasets GSE157896 and GSE55763, with 68.2% and 65.8% of imprinted individuals, respectively (Figure 1, Supplementary Table 2). GSE157896 consists of new-borns whose mothers were Singaporean citizens or permanent residents, with self-reported homogenous Chinese, Indian or Malay ancestry (53), while GSE55763 consists of individuals with Indian Asian ancestry living in the UK (85).

In contrast, datasets consisting primarily of African American individuals (GSE117859 and GSE132203), had the third- and fourth-highest proportions of imprinted individuals - 79.1% and 78.7%, respectively. A third cohort consisting primarily of African American individuals (GSE117860), did not stand out in this regard, with 76.8% of imprinted individuals. As these three datasets consist of other ethnicities in addition to African Americans, we tested whether there is a difference in the proportion of imprinted individuals across ethnic groups. In two of the datasets - GSE117859 and GSE117860, the proportion of imprinted individuals was significantly higher in African Americans than in white individuals (χ2-test p-value < 0.05, Supplementary Figure 2).

The highest proportion of imprinted individuals was observed in datasets GSE121633 and GSE56105, with 83.5% and 79.5% of imprinted individuals respectively. Both GSE121633 and GSE56105 consist of twins. Dataset GSE100227, also consisting of twins, has the fifth-highest percentage of imprinted individuals with 78.7%. However, some datasets consisting of twins had an average proportion of imprinted individuals (74.3% in GSE105018, 75.5% in FTC and 74.5% in SATSA, Figure 1 and Supplementary Table 2).

We found no difference (χ^2^-test p-value > 0.05) in the proportion of imprinted individuals between males and females in 26 datasets (Supplementary Table 2). The only exception was the dataset GSE82273, where the proportion of imprinted individuals was 73.3% (total n=505) in males and 80.7% (total n=384) in females (χ^2^-test p-value = 0.009). This dataset consists of individuals born in Norway with a facial cleft, and unaffected controls matched for the time of birth (33,86). In addition, we found no statistically significant differences in the proportion of imprinted individuals in any of the case-control settings reported (χ^2^-test p-value > 0.05, Supplementary Table 2) and no bias was seen according to the array type utilized (EPIC vs 450K).

### The methylation of nc886 in MZ and DZ twin pairs

We investigated the concordance of the *nc886* methylation level and status between twin pairs. In five datasets, almost all MZ pairs were concordant regarding the *nc886* locus methylation level, whereas a large proportion of DZ twins were discordant for the *nc886* locus methylation level (Table 1, Supplementary Figures 3-6).

**Table 1.**
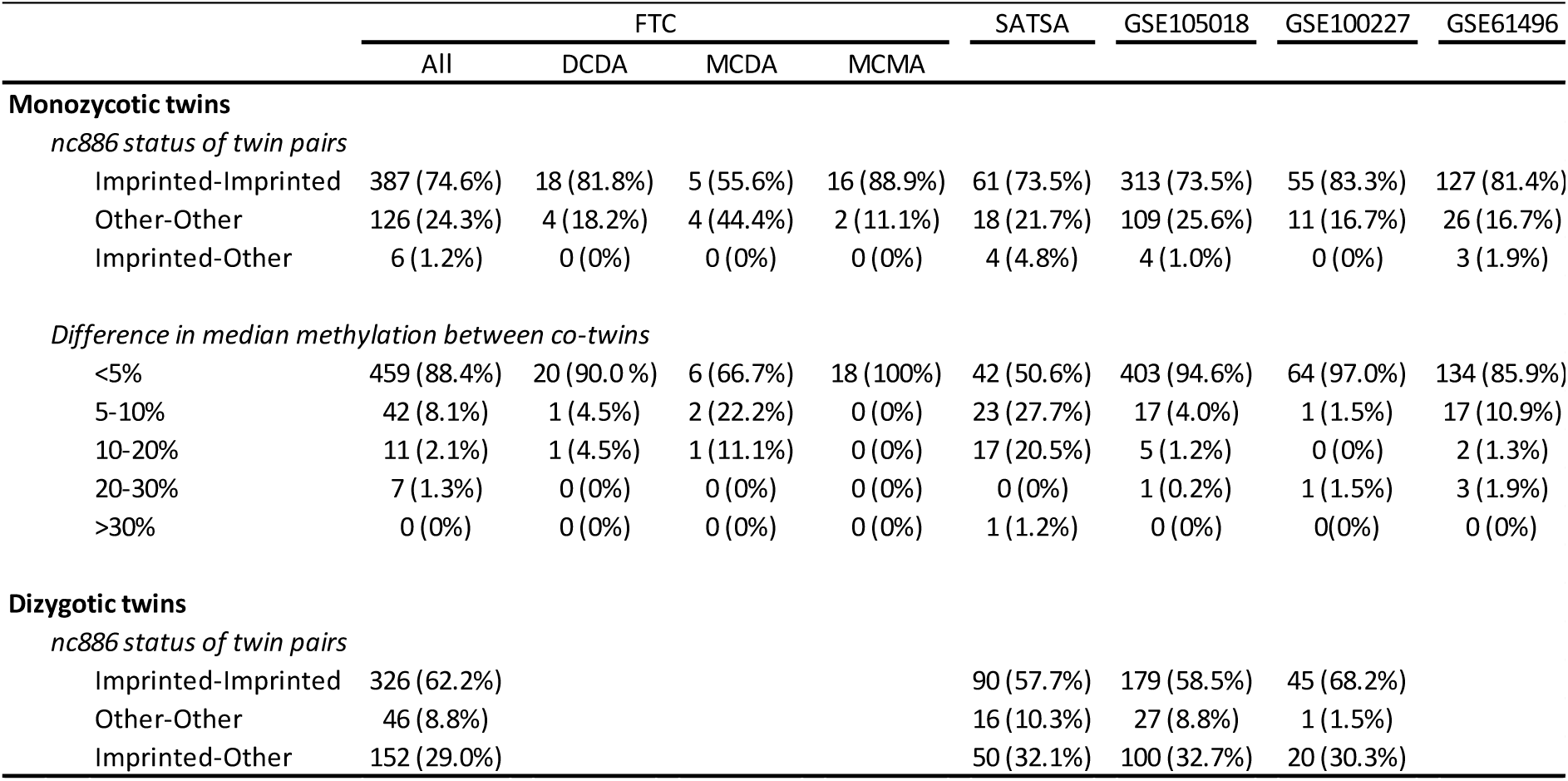
Number of twin pairs discordant for *nc886* methylation status and the absolute difference in the methylation level across twin pairs.

Of the total of 1250 MZ twin pairs investigated, we identified 17 pairs (1.3%), that were clustered to different *nc886* methylation status groups, when individuals were classified as imprinted and other (Table 1). For the datasets in which we could identify the intermediately methylated individuals (GSE61496, GSE105018), we were able to refine the analysis and to include the intermediately methylated and non-methylated groups as well. When considering all three *nc886* status groups (imprinted, intermediately methylated, non-methylated), of the total of 582 MZ pairs investigated, 13 (2.2%) were clustered to different *nc886* status groups. Of these discordant pairs, one co-twin was intermediately methylated while the other co-twin was either imprinted or non-methylated in all cases, i.e. we identified no twin pairs where one co-twin was imprinted while the other was non-methylated (Supplementary Table 3).

Across all twin pair datasets, the absolute difference in the *nc886* methylation level between MZ co-twins was below 0.05 for 88.2% of the pairs. Only 1.0% of pairs had a methylation beta value difference larger than 0.20. Only one MZ pair across all datasets presented a methylation beta value difference over 0.30 (Table 1). For this pair, the methylation beta values for *nc886* locus were 0.38 and 0.71, suggesting that one twin was imprinted and the other had gained methylation also in the paternal allele of *nc886*. These results are in line with our finding that there were no imprinted/non-methylated MZ twin pairs.

We also investigated the absolute difference in the *nc886* methylation level in MZ twin pairs for whom we had information on chorionicity and amnionicity. In DCDA (separated between days 1-3 after fertilization, 22 pairs) and MCDA (separated after day 3, but prior to implantation, 9 pairs) twin pairs, we observed that in 64.5% of the twin pairs the within-pair difference in their median methylation beta values was below 0.025. For the remaining twin pairs, the within-pair difference was 0.025-0.05 in 19.4% of the pairs and above 0.05 in 16.1% of the pairs. In contrast, in MCMA twin pairs (18 pairs), which are separated only after implantation, the within-pair difference in median methylation value was below 0.025 for all pairs.

Four of the five twin cohorts also contained DZ twin pairs. Of these, 30.3%, 35.0%, 29.0%, and 32.1% were *nc886* methylation status (imprinted/other) discordant pairs (Table 1). Given the proportions of the *nc886* methylation status groups in these four datasets, with random pairings, the expected proportion of discordant pairs would be 39.8%, 37.7%, 35.7%, and 38.8%, respectively (details on proportions and expected proportions in Supplementary Table 4). For all four datasets, the proportion of identified discordant pairs was lower than expected, and for FTC, the largest of the available cohorts, the difference was statistically significant (29.0% vs 35.7%, χ2-test p-value = 0.02).

### The methylation level of nc886 differs in the cerebellum and in skeletal muscle

Dataset GSE64491 consists of DNA methylation data for 30 tissues from a 112-year-old female, with tissues analysed in duplicates or quadruplicates. As seen in Figure 2 and Supplementary Figure 7, the methylation level of the *nc886* locus was higher in the cerebellum as compared to other tissues, with the methylation beta value above 0.70 for most probes in this locus. In addition, muscle and diaphragm showed slightly higher methylation beta values as compared to other tissues. For other tissues in dataset GSE64491, variation in the level of methylation at the *nc886* locus between tissues was similar in magnitude to what can be observed between the replicates in the data (Supplementary Table 5). Interestingly, unlike skeletal muscle and diaphragm, the *nc886* methylation level of heart was not elevated, but was comparable to other tissues in the dataset (Supplementary Figure 7). As a reference, we examined the methylation level of six known imprinted genes and did not observe a marked difference in the methylation level between the cerebellum or muscle and other tissues (Supplementary Figure 8). For these six genes, variation in the methylation level between tissues and between the replicates was smaller as compared to the *nc886* locus (Supplementary Table 5).

**Figure 2.**
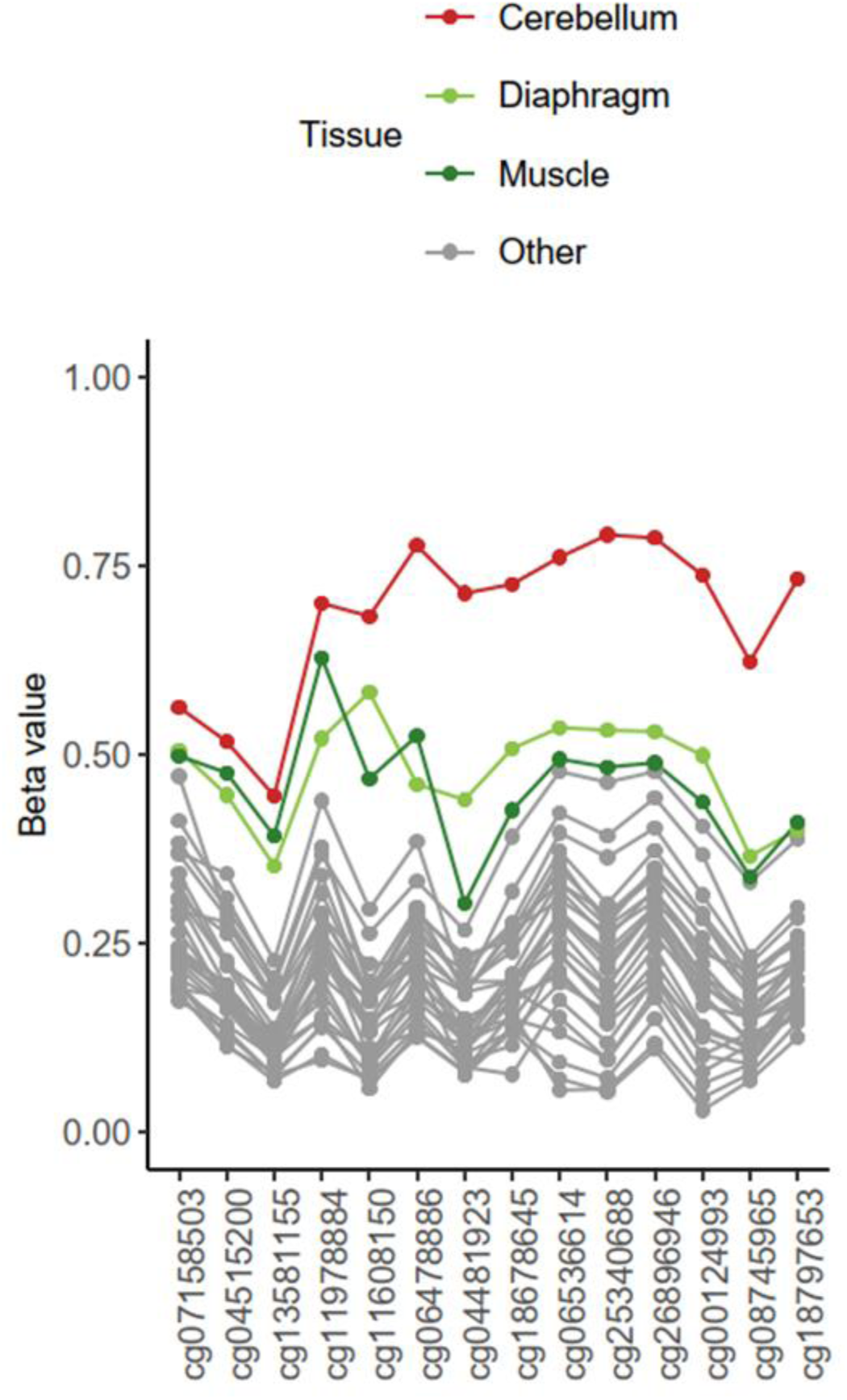
Methylation level of *nc886* locus in 30 tissues of a 112 year-old woman. Cerebellum has considerably higher level of methylation as compared to other tissues. Muscle and diaphragm also show slightly elevated levels of methylation as compared to other tissues. For a figure with all tissues presented in colour, see Supplementary Figure 7.

To verify the methylation level of the *nc886* locus in the cerebellum, dataset GSE134379, containing methylation data from the cerebellum and middle temporal gyrus (MTG), and GSE72778, containing data from the cerebellum and five other brain regions (frontal lobe, hippocampus, midbrain, occipital lobe and temporal lobe) were investigated. In both data-sets, regions of the cerebrum and midbrain present a bimodal *nc886* methylation pattern, while in the cerebellum, the median methylation level follows a unimodal distribution and is higher, close to 0.70 (Figure 3, Supplementary Figure 9).

**Figure 3.**
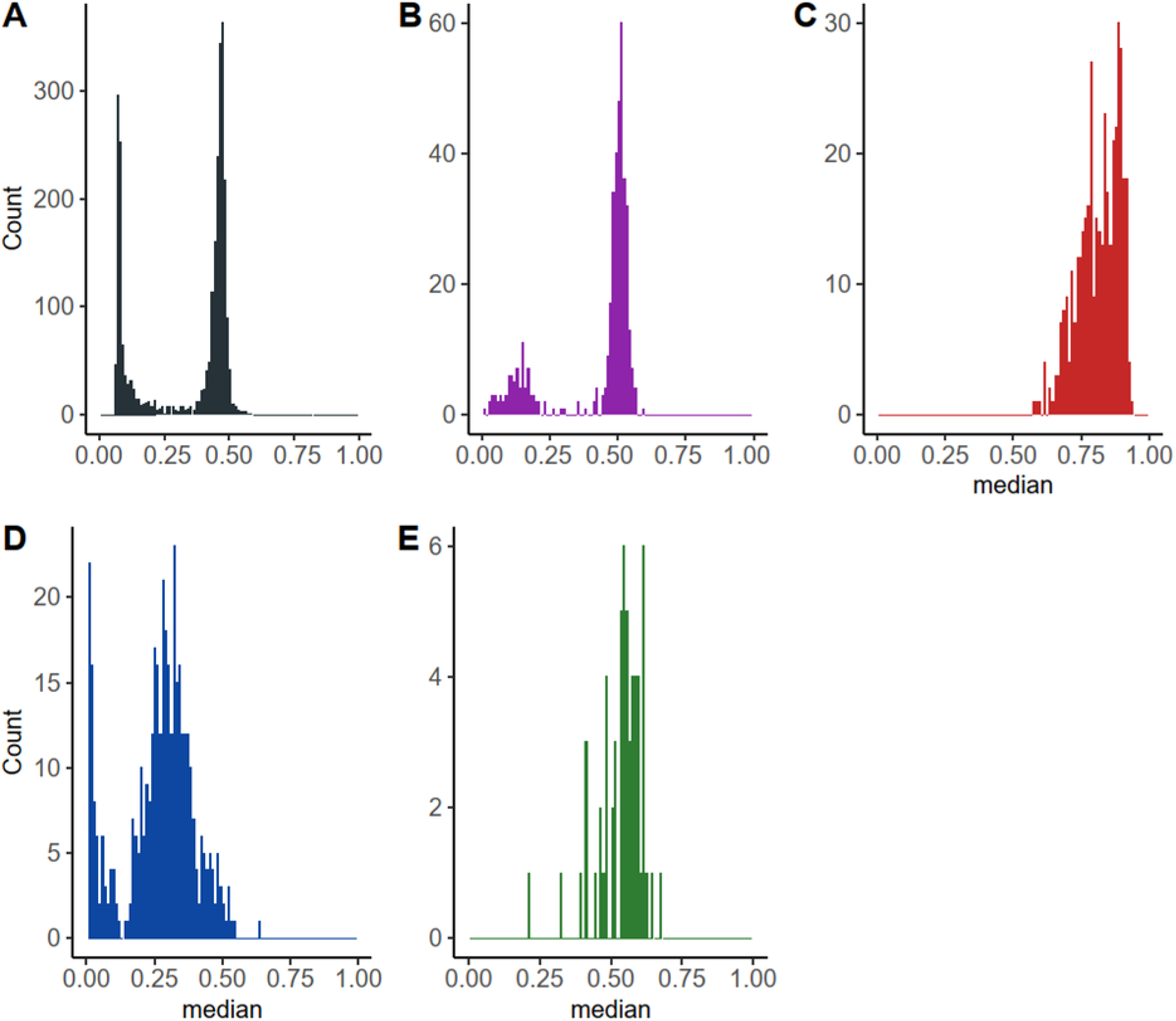
Histograms of *nc886* locus methylation median in different tissues. A) Blood, GSE55763, n = 2664, B) middle temporal gyrus (MTG), GSE134379, n=404, C) cerebellum, GSE134379, n= 404 (same individuals as in B), D) placenta, GSE167885, n = 411, and E) muscle, GSE61454, n = 60. As compared to blood and MTG, cerebellum shows a unimodal distribution with elevated methylation levels in *nc886* locus. Also in muscle (E) *nc886* methylation showed a unimodal distribution. In placenta the methylation level at *nc886* locus presented a bimodal distribution, but the overall methylation level was lower as compared to blood.

The methylation level of the *nc886* locus in the cerebellum was high both in individuals with an imprinted and a non-methylated *nc886* (clustered according to the methylation levels in the cerebrum or MTG) in both datasets studied (Figure 4 and Supplementary Figure 10). Despite the unimodal distribution of the *nc886* methylation level in the cerebellum, we observed a difference in the methylation level of the *nc886* locus in the cerebellum between imprinted and non-methylated individuals (p-value < 0.001 in both datasets). The six known imprinted genes analysed did not present differences in the methylation level between the cerebellum and other brain regions (Supplementary Figure 11).

**Figure 4.**
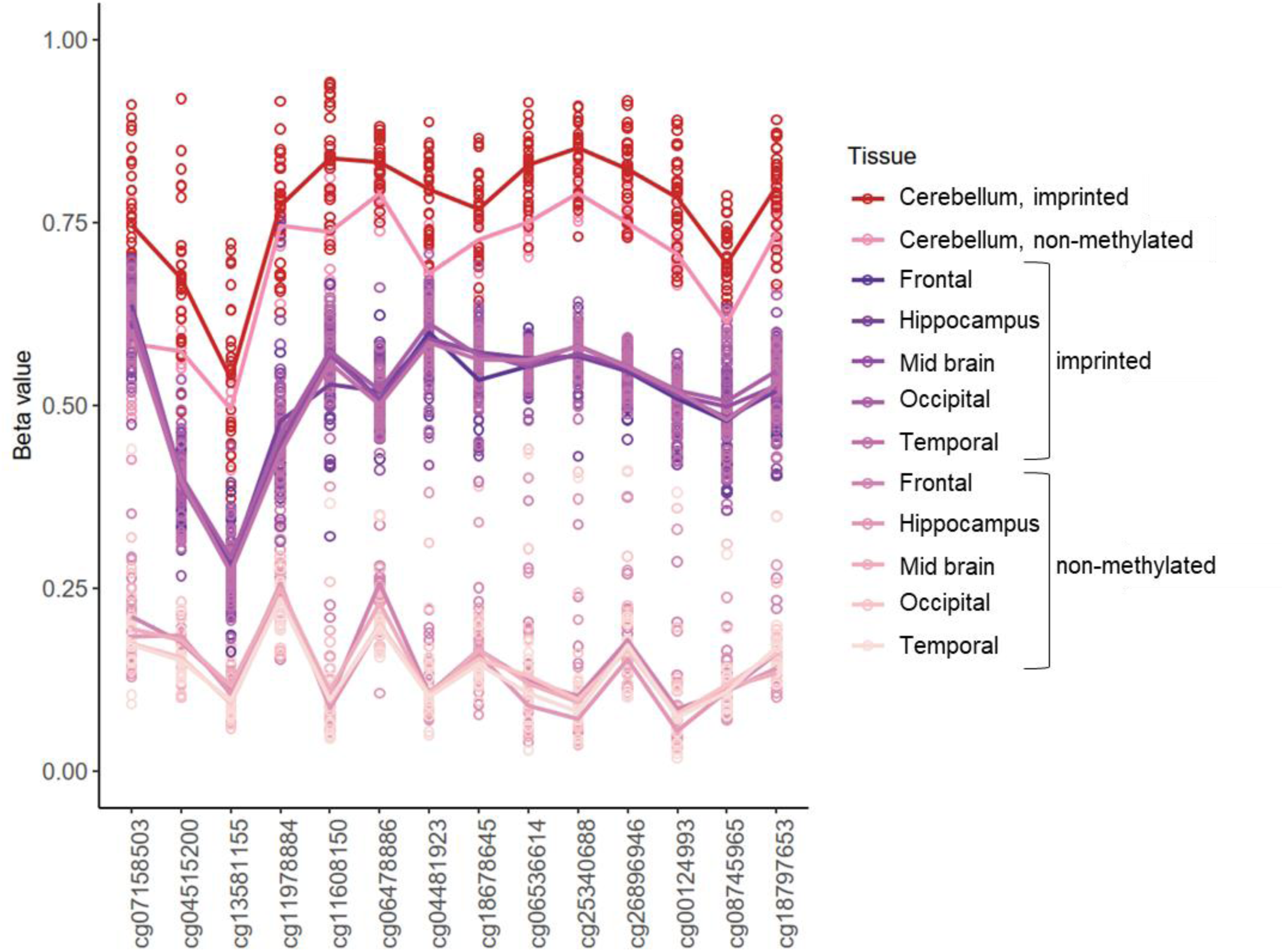
Methylation level of *nc886* in cerebellum and in five other brain regions (GSE72778). Individuals have been grouped to imprinted and non-methylated based on data of the five brain regions where *nc886* displayed a bimodal methylation pattern (frontal lobe, hippocampus, mid brain, occipital lobe and temporal lobe). As compared to other brain regions, cerebellum shows higher levels of methylation in both groups, but there is a statistically significant difference in the methylation between individuals clustered as imprinted and non-methylated. Similar pattern can be observed in dataset GSE134379 (Supplementary Figure 10).

To further investigate the methylation level of the *nc886* locus in muscle, we investigated six additional datasets, three of which consisted only of muscle samples (GSE142141, GSE151407, GSE171140), and three for which other tissues were also available (GSE61454 - skeletal muscle, visceral and subcutaneous fat, as well as liver; ERMA - muscle tissue and blood; FTC - muscle tissue, adipose tissue, and blood). In all of the datasets, the methylation level of the *nc886* locus in muscle presented a unimodal distribution, centred slightly above 0.50 (Figure 3, Supplementary Figures 12-14). In GSE61454, ERMA and FTC, other tissues presented the expected bimodal methylation distribution at the *nc886* locus (Supplementary Figures 13 and 14). Despite the unimodal methylation level observed in muscle, there was a difference in the methylation level at the *nc886* locus in muscle between imprinted and non-methylated individuals (as clustered based on the *nc886* methylation levels in other tissues) in FTC and GSE61454 (Mann-Whitney U-test p-value < 0.001) (Figure 5, Supplementary Figure 15). Moreover, in FTC, the median methylation levels correlated well across all three tissues (adipose tissue, muscle, blood; p<2*10^-12^). Individuals presenting intermediately methylated *nc886* in blood also had intermediate methylation levels in other tissues (Supplementary Figure 16).

**Figure 5.**
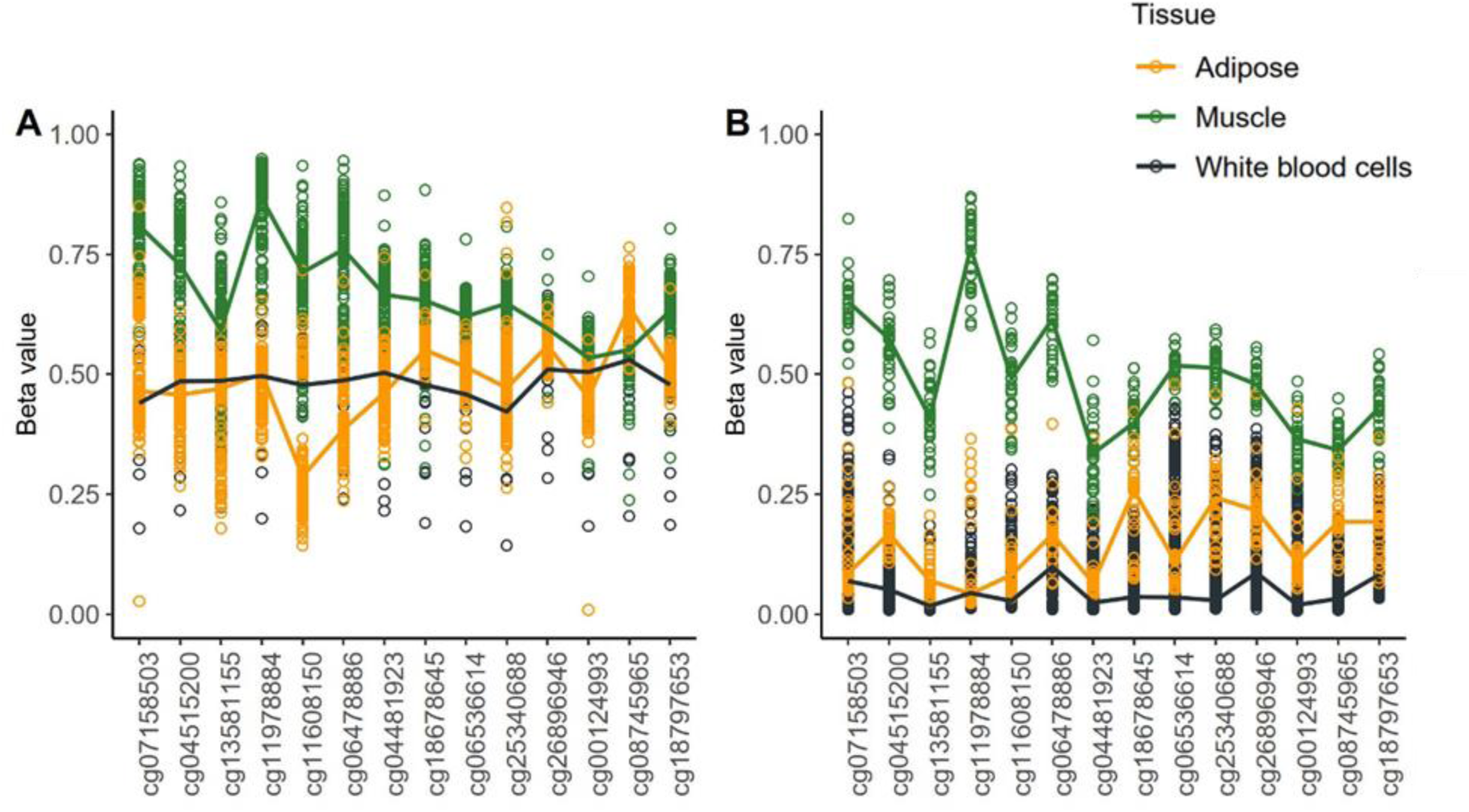
Methylation level of *nc886* locus in FTC in A) imprinted individuals (as clustered based on *nc886* methylation levels in blood) and B) non-methylated individuals. Despite the unimodal methylation level in muscle (Supplementary Figure 13), we observed a difference in the methylation level of *nc886* locus in muscle between imprinted and non-methylated individuals.

We also investigated other available datasets for potential atypical methylation patterns in the *nc886* locus. Skin (GSE90124) and buccal swabs (NELLI, GSE128821) displayed the expected bimodal distribution and the expected level of methylation at the *nc886* locus (Supplementary Figure 17). For the placenta (Figure 3, Supplementary Figures 18 and 19), we observed a slight downward shift in the methylation level at the *nc886* locus in four datasets (GSE167885, GSE75248, GSE71678, GSE115508). While the methylation pattern in the placenta followed a bimodal distribution, a clustering analysis could not establish a clear division between imprinted and non-methylated individuals, as compared to the majority of other tissues studied. In dataset GSE115508, we did not observe a corresponding downward shift in the methylation values in amnion or chorion (Supplementary Figure 19). Similarly, foetal cord tissue (GSE157896) did not display a downward shift in the *nc886* methylation level (Supplementary Figure 1).

In sperm, we observed a methylation level close to 0 at the *nc886* locus, as is expected for a maternally imprinted gene (Supplementary Figure 20). In addition to methylation data from sperm, dataset GSE149318 also contained methylation data from blood of the same individuals. We did not observe a difference in the sperm *nc886* methylation level between blood-derived imprinted and non-methylated individuals (Mann-Whitney U-test p-value > 0.05).

## Discussion

Here, we confirm that the proportion of individuals with an imprinted *nc886* locus is approximately 75% in majority of populations and show in 32 datasets from variable historic and geographic origins, that the variation in this proportion is very limited, especially in populations consisting of white singletons. More varying proportions can be observed in populations consisting of other ethnicities, and in twins. We show that in MZ twin pairs, the methylation level of the *nc886* locus is highly similar, especially in MCMA twin pairs. Finally, we confirm that the *nc886* methylation pattern is stable in the majority of somatic tissues, but also describe two exceptions – the cerebellum and skeletal muscle. These findings allow us to refine the hypotheses on timing and determinants of the polymorphically imprinted *nc886* and how the methylation in this locus varies in tissues and individuals originating from the same zygote.

### Variation of the nc886 methylation status group proportions is limited across populations

It has previously been reported that, based on individual cohorts, the proportion of individuals with an imprinted *nc886* locus is approximately 75%, with the remaining 25% presenting a non-methylated *nc886* locus (9–11). Here, utilizing 32 cohorts and over 30 000 individuals, we can confirm that the average proportion of imprinted individuals is 75%. Especially in white singletons, the variation in the proportion of individuals with an imprinted *nc886* locus is limited. The lowest proportion of individuals with an imprinted *nc886* is observed in cohorts of East Asian origin, while the highest proportion of individuals with an imprinted *nc886* is observed in cohorts consisting of African American individuals or twins. Previously, in a cohort of 82 Korean women, the proportion of individuals with methylation levels indicating an imprinted *nc886* locus was reported to be 65.9% (87), comparable to our findings here (Figure 1, Supplementary Table 2). We, and others, have previously shown that genetic variation is not associated with the establishment of the *nc886* methylation pattern (10,11,18), but, as these results are based on mainly white populations, we can’t rule out the possibility that other ethnic groups would present genetic variation that would affect the establishment of the *nc886* imprint. Another explanation for these findings is that the lifestyle or environmental conditions of these populations are affecting the proportions of imprinted individuals. However, dataset GSE157896 consists of individuals born in Singapore (53) and dataset GSE55763 consists of individuals of East Asian origin, some of whom have been born in the UK and others in Asia (29), suggesting that, at least in this case, shared genetics, rather than shared environmental conditions, may affect the proportion of imprinted individuals. Our results again highlight the need to include more diverse populations in genetic association studies (88).

### The methylation status of nc886 is not associated with sex

We identified no difference in the proportion of imprinted individuals between males and females, with the exception of one dataset - GSE82273. This dataset consists of individuals born with a facial cleft and unaffected controls matched for the time of birth (33,86). In a previous study, facial clefts were associated with hypomethylation of the *nc886* region (89). However, in this study, the *nc886* methylation was analysed as a continuous variable instead of categorical. The prevalence of facial clefts has also been reported to vary according to sex, whether it is female- or male-biased depending on the type of the cleft (90). Unfortunately, the information on the case/control status was not included in dataset GSE82273, and thus we were unable to test whether the observed difference in proportions is due to the sex bias of facial clefts. As we observed no difference in the proportion of imprinted individuals between males and females in 26 other datasets (in a total of 27 362 individuals), we assume that the observed difference in this one dataset is due to the bias caused by the case/control setting of the dataset and conclude that the *nc886* methylation status is not associated with sex.

### nc886 imprint is stable across majority of somatic tissues

Previous studies have shown that the *nc886* methylation status is stable within one individual in all tissues analysed (10,16), and we have previously shown this in different blood cell subtypes (11). Here, we confirm these findings in over 30 somatic tissues but also report exceptions to this pattern, namely cerebellum and skeletal muscle. In both cerebellum and skeletal muscle, visual inspection revealed a unimodal DNA methylation pattern at the *nc886* locus, with a methylation level of approximately 0.70 in the cerebellum and approximately 0.50 in skeletal muscle. Despite the unimodal methylation pattern in these tissues, there was a slight difference in the methylation level of *nc886* between individuals who were imprinted and non-methylated (according to clustering analysis on other tissues). This suggests that the methylation pattern established in early development in these tissues is not completely reset and established anew, but that the methylation level is built upon the existing methylation state. This is further corroborated by our finding that the methylation level of *nc886* in muscle is strongly correlated with the methylation level in other somatic tissues. It is not known what mechanism is responsible for this increase in methylation at the *nc886* locus in these tissues. In contrast to the cerebellum and skeletal muscle, in the placenta, the methylation level of the *nc886* locus was slightly decreased as compared to other analysed tissues. Placenta has been previously described to have aberrant profiles of imprinted genes and also present a multitude of secondary differentially methylated regions (91). In line with previous studies (9,13), we also show here that the *nc886* locus is non-methylated in sperm and see no difference in the methylation levels between men who present either imprinted or non-methylated *nc886* locus in other tissues.

We, and others, have previously shown that the *nc886* methylation is tightly associated with the level of nc886 RNAs (11,12,92), with the imprinted individuals having lower levels of these RNAs as compared to the non-methylated individuals. Therefore, we can speculate that the cerebellum and skeletal muscle have lower levels of these RNAs as compared to other tissues, while the placenta has higher levels of these RNAs. As the function of these RNAs is not known (14,15), different regulation patterns in these specific tissues might offer possibilities to further hypothesize their role.

In addition to being stable across tissues, the methylation level of *nc886* has been shown to be stable through follow-up, from childhood to adolescence (16) and from adolescence to adulthood (11). However, in granulosa cells, the methylation level of the *nc886* locus has been shown to be affected by age, with women over the age of 40 showing higher methylation values as compared to women under the age of 30 (93). Granulosa cells are the somatic cell compartment in the follicle and are crucial for oogenesis (94,95). As it has been previously shown that maternal age is associated with the *nc886* methylation status of the offspring (9,11), it is interesting to speculate whether the altered methylation status of the granulosa cells is associated with this phenomenon.

### Establishment of nc886 imprint

Current literature suggests that periconceptional conditions affect the establishment of the *nc886* methylation pattern (9–11,16) and that the imprinting of *nc886* is an early embryonic event happening between days four and six after fertilization (13). This notion is in slight conflict with the finding that the variation in the proportion of imprinted individuals is limited across populations. If periconceptional conditions would have a substantial role in the establishment of the methylation of the *nc886* locus, one would expect to see more differences between cohorts from different countries or between different birth cohorts and, in contrast, fewer differences between DZ twin pairs. For example, we see only minor differences in the prevalence of imprinted individuals in Finnish cohorts born in the 1960s and 1970s (YFS; 73.5%), the 1950s through 1980s (FTC;75.7%) or in 2007/2008 (NELLI; 74.1%), even though the nutritional status of Finnish expecting mothers has drastically changed during this time (96,97). Furthermore, if the non-methylated status would be caused by adverse pregnancy conditions, such as lack of energy or certain nutrients, one could expect that the proportion of non-methylated individuals would be significantly decreased in populations with good nutritional status. For example, if the lack of folate would be causal in the establishment of the non-methylated *nc886* methylation pattern (16,98), the number of these individuals should have been more drastically diminished in cohorts born after the folate supplementation recommendations (97,99). For DZ twins, who share the pregnancy but originate from different zygotes, we showed that approximately 1/3 of pairs are discordant regarding the *nc886* methylation status. While this was slightly fewer pairs than would be expected by chance, the difference was subtle and statistically significant only in one dataset studied.

Another plausible hypothesis explaining the shown associations between *nc886* and periconceptional conditions is that the methylation status is determined already in the oocyte, as suggested also by Carpenter et al. (10), and that the slight variation observed in the *nc886* status proportions at the population level is due to either epigenotype offering a survival advantage in specific pregnancy conditions. The establishment of the *nc886* methylation status already in the oocyte is also supported by the fact that in line with results by Carpenter et al. (9), we observed no MZ twin pairs with one co-twin being imprinted and the other non-methylated. This is also true in the subgroup of twins from FTC, who have been reported to be dichorionic, and thus have been separated 1-3 days after conception, indicating that the process leading to either non-methylated or imprinted *nc886* loci was completed before this time point.

Our results also suggest that the process that leads to individuals presenting intermediate *nc886* methylation pattern is over before implantation. In MZ twins reported to be separated after implantation, the median methylation in *nc886* is nearly identical (difference less than 2.5%). Furthermore, individuals who present intermediate methylation levels in their blood also present very similar levels in their adipose tissue, in line with previous findings in different blood cell populations (11), suggesting stability of the methylation level, including the intermediate state, through tissue differentiation. When taken together with the temporal stability of also the intermediate methylation levels (11), we can suggest that the ratio of cells with an imprinted or a non-methylated maternal allele of *nc886* in individuals presenting intermediately methylated *nc886* status is established before implantation, concurrently to the global de- and re-methylation waves in the embryo (100) and is then reflected on the individual for the rest of their life.

Previous reports on the association of periconceptional conditions and the *nc886* methylation status (9–11,16) could be explained by selective survival in certain pregnancy conditions, instead of these conditions directly affecting the establishment of the methylation status. As shown by our results, an example of pregnancy conditions where certain *nc886* status might be advantageous/disadvantageous is twin pregnancy. In the population cohorts studied, twin cohorts had high proportions of imprinted individuals, and in DZ twins the number of pairs with discordant *nc886* methylation status was slightly lower than would be expected by chance. This suggests that twin pregnancy might be favourable to foetuses with imprinted *nc886* loci or, in the case of DZ twins, to pairs with concordant *nc886* methylation status.

### Limitations of the study

Our results are descriptive in nature, and thus need to be interpreted as hypothesis-generating rather than conclusive. Clustering of individuals into *nc886* methylation status groups is, to some extent, affected by different data pre-processing methods, but we have tried to mitigate this by carefully comparing different pre-processing methods. The paucity of datasets consisting of individuals of multiple ethnicities limits our possibilities to draw firm conclusions on the effect of ethnicity on *nc886* methylation status proportions.

### Conclusion

Current literature suggests that the polymorphic imprinting of *nc886* is not due to genetic variation in white populations, but as our results show more variation in the proportion of individuals with an imprinted *nc886* in non-white cohorts, the genetic analyses should be repeated in more diverse populations. Based on our results and current literature, we hypothesize that DNA methylation of the *nc886* locus is established in the growing oocyte and that the variation in the proportion of imprinted individuals in a population could be due to survival advantage/disadvantage in certain pregnancy conditions, illustrated in Supplementary Figure 21. After implantation, the methylation level of *nc886* is preserved across studied somatic tissues, with the exception of cerebellum and skeletal muscle. In all individuals, *nc886* locus gains methylation in these tissues, even though the methylation levels still associate with the *nc886* status established earlier in the development.

## Summary points

- Variation in the proportion of individuals with an imprinted *nc886* is very modest, with approximately 75% of individuals being imprinted across populations
- The observed variation is mainly limited to non-white ethnic groups and twin cohorts
- Methylation status of *nc886* was not associated with sex or any of the case/control setting investigated
- Methylation level of *nc886* is increased in cerebellum and in skeletal muscle, but is uniform in other somatic tissues
- Placenta presents lower methylation levels than majority of somatic tissues, but binomial methylation pattern can still be detected
- Monozygotic twins show highly similar *nc886* methylation levels, which is even more pronounced on twins separated at a later date in development
- Approximately 30% of dizygotic twin pairs are discordant for *nc886* methylation status
- We suggest that methylation status of *nc886* is established in the oocyte, and that the slight variation observed across populations could be due to selective survival advantage of the foetus in certain conditions

## Reference annotations

**of considerable interest

** Carpenter BL, Zhou W, Madaj Z, DeWitt AK, Ross JP, Grønbæk K, et al. Mother-child transmission of epigenetic information by tunable polymorphic imprinting. Proc Natl Acad Sci U S A. 2018 Dec 18;115(51):E11970–7.

*Describes how methylation status of nc886 is associated with prenatal environment, and that the methylation status is associated with the offspring’s phenotype*.

** Carpenter BL, Remba TK, Thomas SL, Madaj Z, Brink L, Tiedemann RL, et al. Oocyte age and preconceptual alcohol use are highly correlated with epigenetic imprinting of a noncoding RNA (nc886). Proc Natl Acad Sci U S A. 2021 Mar 23;118(12):e2026580118.

*Further confirms the association between periconceptional conditions and offspring’s nc886 methylation status and describes that 75% of oocytes also present imprinted nc886 locus*.

** Silver MJ, Kessler NJ, Hennig BJ, Dominguez-Salas P, Laritsky E, Baker MS, et al. Independent genomewide screens identify the tumor suppressor VTRNA2-1 as a human epiallele responsive to periconceptional environment. Genome Biology. 2015 Jun 11;16(1):118.

*Identifies nc886 as metastable epiallele, that is responsive to environmental conditions*.

** Marttila S, Viiri LE, Mishra PP, Kühnel B, Matias-Garcia PR, Lyytikäinen LP, et al. Methylation status of nc886 epiallele reflects periconceptional conditions and is associated with glucose metabolism through nc886 RNAs. Clin Epigenetics. 2021 Jul 22;13(1):143.

*Further confirms the association between periconceptional conditions and offspring’s nc886 methylation status, describes the association with offspring’s phenotype. Describes the association of nc886 methylation and nc886 RNA levels*.

*of interest

* Kostiniuk D, Tamminen H, Mishra PP, Marttila S, Raitoharju E. Methylation pattern of polymorphically imprinted nc886 is not conserved across mammalia. PLOS ONE. 2022 Mar 16;17(3):e0261481.

*Characterises nc886 methylation in primates, indicating that the locus is not polymorphically imprinted in other species than in humans*

* Romanelli V, Nakabayashi K, Vizoso M, Moran S, Iglesias-Platas I, Sugahara N, et al. Variable maternal methylation overlapping the nc886/vtRNA2-1 locus is locked between hypermethylated repeats and is frequently altered in cancer. Epigenetics. 2014 May 6;9(5):783–90.

*Describes how nc886 is a metastable epiallele, the establishment of which is an early embryonic event and shows that nc886 is maternally imprinted*.

* Treppendahl MB, Qiu X, Søgaard A, Yang X, Nandrup-Bus C, Hother C, et al. Allelic methylation levels of the noncoding VTRNA2-1 located on chromosome 5q31.1 predict outcome in AML. Blood. 2012 Jan 5;119(1):206–16.

*Shows that in population 75% of individuals present imprinted nc886 loci, and that the expression of nc886 RNAs are associated with the methylation level in the locus*.

## Ethical declarations

YFS: approved by the 1st ethical committee of the Hospital District of Southwest Finland on September 21st, 2010, and by local ethical committees (1st Ethical Committee of the Hospital District of Southwest Finland, Regional Ethics Committee of the Expert Responsibility area of Tampere University Hospital, Helsinki University Hospital Ethical Committee of Medicine, Research Ethics Committee of the Northern Savo Hospital District, and Ethics Committee of the Northern Ostrobothnia Hospital District).

NELLI: This follow-up study and written informed consent procedure were approved by the medical ethics committees of the Pirkanmaa hospital district (R14039)

SATSA: study was approved by the ethics committee at Karolinska Institutet with Dnr 2015/1729-31/5.

LURIC: The study plan was approved by the ethics committee of the State Chamber of Physicians of Rhineland-Palatinate.

KORA: All study methods were approved by the ethics committee of the Bavarian Chamber of Physicians, Munich.

DILGOM: original FINRISK study has been approved by Coordinating Ethics Committee of the HUS Hospital District, decision numbers 229/E0/2006 and 332/13/03/00/2013. FINRISK and DILGOM study materials have been transferred to THL Biobank in accordance with the notification procedure permitted by the Finnish Biobank Act.

FTC: Data collection and analysis were approved by the ethics committee of the Helsinki University Central Hospital (Dnro 249/E5/01, 270/13/03/01/2008, 154/13/03/00/2011).

ERMA: study was approved by the ethics committee of the Central Finland health care district in 2014 (K-SSHP Dnro 8U/2014).

## Acknowledgements

The authors wish to thank all the researchers who have participated in the collection of the utilized datasets and made their data publicly available. The DILGOM data used for the research were obtained from THL Biobank (study number THLBB2021_22). We thank all study participants for their generous participation at THL Biobank and at National FINRISK and DILGOM Studies. We also thank Daria Kostiniuk (BSc) for the in-depth discussions of the establishment of the *nc886* methylation pattern.

