## Supplemental data 1 for "Methylation status of *VTRNA2-1*/*nc886* is stable across human populations, monozygotic twin pairs and in majority of somatic tissues"

**Supplementary Data 1**

### Methylation status of *VTRNA2-1*/*nc886* is stable across human populations, monozygotic twin pairs and in majority of somatic tissues

Saara Marttila, Hely Tamminen, Sonja Rajić, Pashupati P Mishra, Terho Lehtimäki, Olli Raitakari, Mika Kähönen, Laura Kananen, Juulia Jylhävä, Sara Hägg, Thomas Delerue, Annette Peters, Melanie Waldenberger, Marcus E Kleber, Winfried März, Riitta Luoto, Jani Raitanen, Elina Sillanpää, Eija K Laakkonen, Aino Heikkinen, Miina Ollikainen, Emma Raitoharju

#### *Clustering of individuals based on nc886 methylation status*

As all datasets in GEO are not available as raw data, we wanted to establish a reproducible way, irrespective of normalization method used, to cluster the individuals to *nc886* methylation status groups. DNA methylation data GSE157896 (1), available in GEO (2) was downloaded as raw idat-files and extracted with minfi package function read.metharray.exp. Four normalization methods with default settings were tested: SWAN from both minfi (SWAN M) and wateRmelon (SWAN W), quantile normalization from minfi as well as dasen from wateRmelon (3–5). In addition, raw beta values were extracted by preprocessRaw-function from minfi package.

The differentially normalized datasets, along with raw data, were clustered with either hierarchical or k-means clustering to 2, 3 and 4 groups. For each cluster, the methylation median of *nc886* locus was calculated. Clusters where median beta value of *nc886* locus was >0.40 were interpreted to be imprinted, clusters where median of *nc886* locus was < 0.15 were interpreted to be non-methylated and clusters where median of *nc886* locus was 0.15-0.40 were interpreted to be intermediately methylated. For each normalization and clustering method we then compared whether individuals were in the same category (imprinted, intermediately methylated or non-methylated or only imprinted and other, i.e. either intermediately methylated and non-methylated combined) for each data normalization method.

With three groups (imprinted, intermediately methylated and nonmethylated) there were considerable inconsistencies between non-methylated and intermediately methylated groups with both clustering methods, with up to 5% of individuals in different methylation status groups across different normalization methods. With two groups (imprinted and other) and hierarchical clustering, there were still inconsistencies across normalization methods with up to 4% of individuals clustered to different *nc886* methylation status groups. However, with k-means clustering, only 2 (out of 1019, 0.02%) individuals were grouped to different methylation status groups across all methods (Supplementary Data Table SD1).

*Supplementary Data Table SD1. Reproducibility of nc886 status groups across different normalization methods. Dataset GSE157896 was normalized with four different methods, and individuals were clustered to three groups with k-means clustering. From these groups, we identified the imprinted individuals and grouped non-methylated and intermediately methylated as 'other'. Across different normalization methods, only 2 individuals out of 1019 (0.02%) were clustered to different nc886 methylation status groups.*

| | | quantile | | SWAN M | | SWAN W | | raw | |
|---|---|---|---|---|---|---|---|---|---|
| | | imprinted | other | imprinted | other | imprinted | other | imprinted | other |
| dasen | imprinted | 694 | 0 | 694 | 0 | 694 | 0 | 694 | 0 |
| | other | **1** | 324 | **1** | 324 | **1** | 324 | **2** | 323 |
| quantile | imprinted | | | 695 | 0 | 695 | 0 | 695 | 0 |
| | other | | | 0 | 324 | 0 | 324 | **1** | 323 |
| SWAN M | imprinted | | | | | 695 | 0 | 695 | 0 |
| | other | | | | | 0 | 324 | **1** | 323 |
| SWAN W | imprinted | | | | | | | 695 | 0 |
| | other | | | | | | | **1** | 323 |

To further confirm this, we repeated the different normalization methods to dataset GSE125105 (6) and clustered the individuals as described above with k-means clustering to imprinted and ‘other’. In this dataset, no more than 5 (out of 699, 0.72%) individuals were grouped to different *nc886* methylation status groups across all methods (Supplementary Data Table SD2).

*Supplementary Data Table SD2. Reproducibility of nc886 status groups across different normalization methods in GSE125105. Across different normalization methods, only 5 individuals out of 699 (0.72%) were clustered to different nc886 methylation status groups.*

| | | quantile | | SWAN M | | SWAN W | | raw | |
|---|---|---|---|---|---|---|---|---|---|
| | | imprinted | other | imprinted | other | imprinted | other | imprinted | other |
| dasen | imprinted | 515 | **5** | 519 | **1** | 519 | **1** | 519 | **1** |
| | other | 0 | 179 | 0 | 179 | 0 | 179 | 0 | 179 |
| quantile | imprinted | | | 515 | 0 | 515 | 0 | 515 | 0 |
| | other | | | **4** | 180 | **4** | 180 | **4** | 180 |
| SWAN M | imprinted | | | | | 519 | 0 | 519 | 0 |
| | other | | | | | 0 | 180 | 0 | 180 |
| SWAN W | imprinted | | | | | | | 519 | 0 |
| | other | | | | | | | 0 | 180 |

Therefore, all datasets utilized in this study were clustered with k-means clustering to three groups, from which the imprinted clusters (median *nc886* beta value > 0.40) were identified, other clusters for each data set were combined to category ‘other’.
