## Supplementary material for "Methylation status of *VTRNA2-1*/*nc886* is stable across human populations, monozygotic twin pairs and in majority of somatic tissues": Supplemtary Figures 1-21

**Supplementary Figures 1-21**

### Methylation status of *VTRNA2-1/nc886* is stable across human populations, monozygotic twin pairs and in majority of somatic tissues

Saara Marttila, Hely Tamminen, Sonja Rajić, Pashupati P Mishra, Terho Lehtimäki, Olli Raitakari, Mika Kähönen, Laura Kananen, Juulia Jylhävä, Sara Hägg, Thomas Delerue, Annette Peters, Melanie Waldenberger, Marcus E Kleber, Winfried März, Riitta Luoto, Jani Raitanen, Elina Sillanpää, Eija K Laakkonen, Aino Heikkinen, Miina Ollikainen, Emma Raitoharju

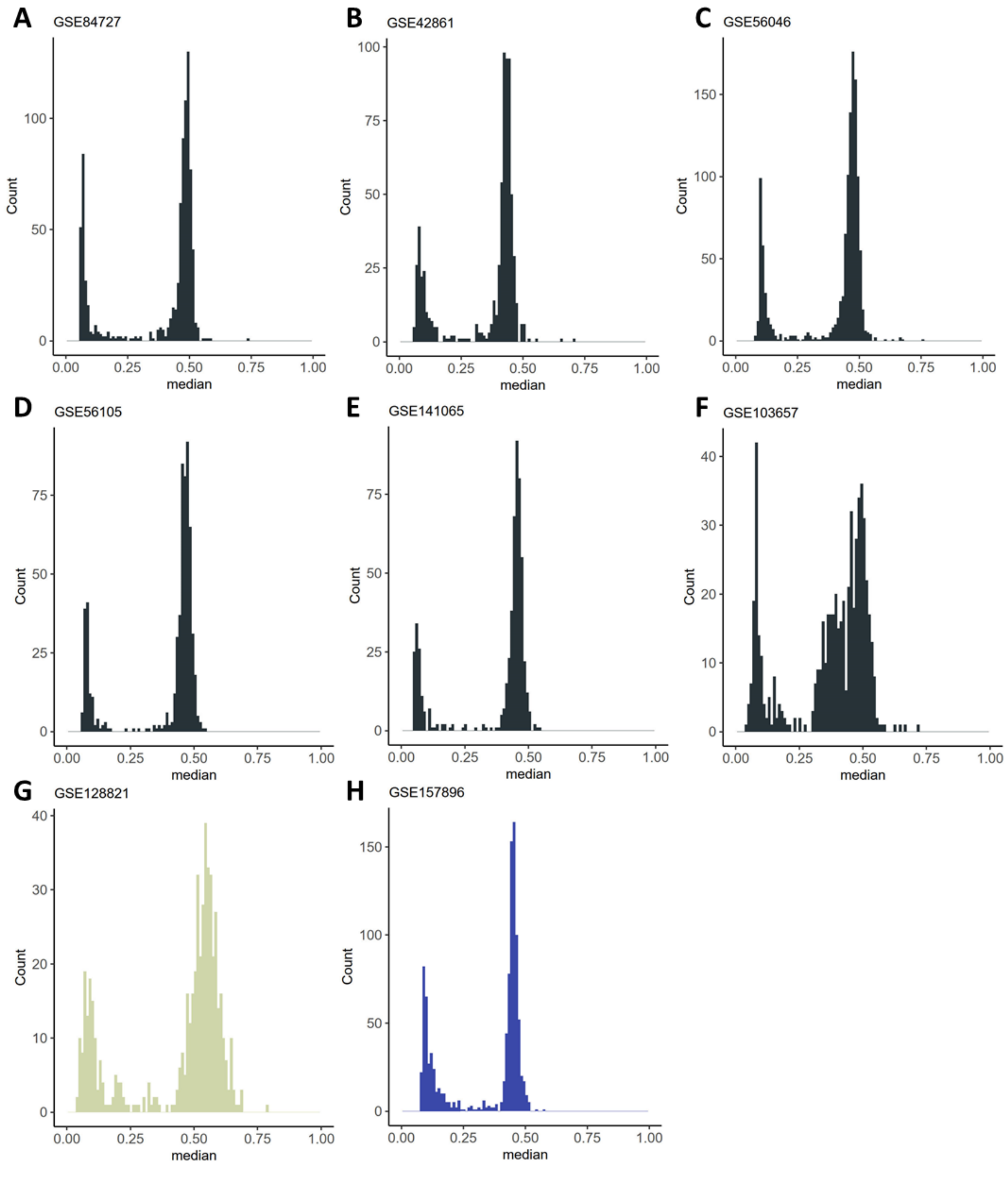


**Supplementary Figure 1.** *A representative example of the median methylation level of nc886 locus in tissues utilized for population cohorts. A) blood (GSE84727), n= 847 B) peripheral blood leukocytes (GSE42861), n=689 C) peripheral blood CD14+ cells (GSE56046), n=1202 D) peripheral blood lymphocytes (GSE56105), n=614 E) umbilical cord blood buffy coat (GSE141065), n=557 F) neonatal blood spot (GSE103657), n=438 G) buccal swab (GSE128821), n=536 H) fetal cord tissue (GSE157896), n=1019. In all tissues we observed the expected bimodal methylation pattern, enabling us to cluster the individuals to nc886 methylation status groups.*

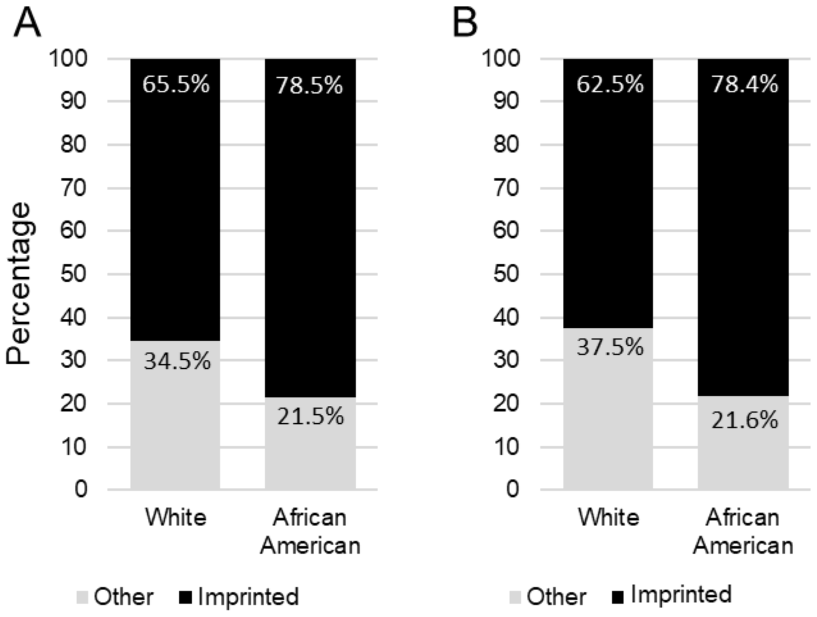


***Supplementary Figure 2.*** *Percentage of individuals identified as imprinted or other in nc886 locus A) in GSE117859 and B) in GSE117860. In both datasets, the difference in the proportion of imprinted individuals was different between white and African American individuals ($\chi$2-test p-value 0.025 and 0.013 respectively). In both datasets, individuals for whom ethnicity was coded as 'other' were also included. In both datasets, we tested whether the proportion of imprinted individuals was different between African American and the combined group of white and 'other'. In GSE117859 the difference was again statistically significant ($\chi$2-test p-value 0.006), but in GSE117860 the difference was not statistically significant ($\chi$2-test p-value 0.067). In these datasets the proportion of imprinted individuals among the white and ethnicity 'other' was notably low (see Figure 1 and Supplementary Table 2 for comparison).*

*Number of individuals, in GSE117859, African American n=522, white n=58, other n=28; in GSE117860, African American n=426, white n=48, other n=55.*

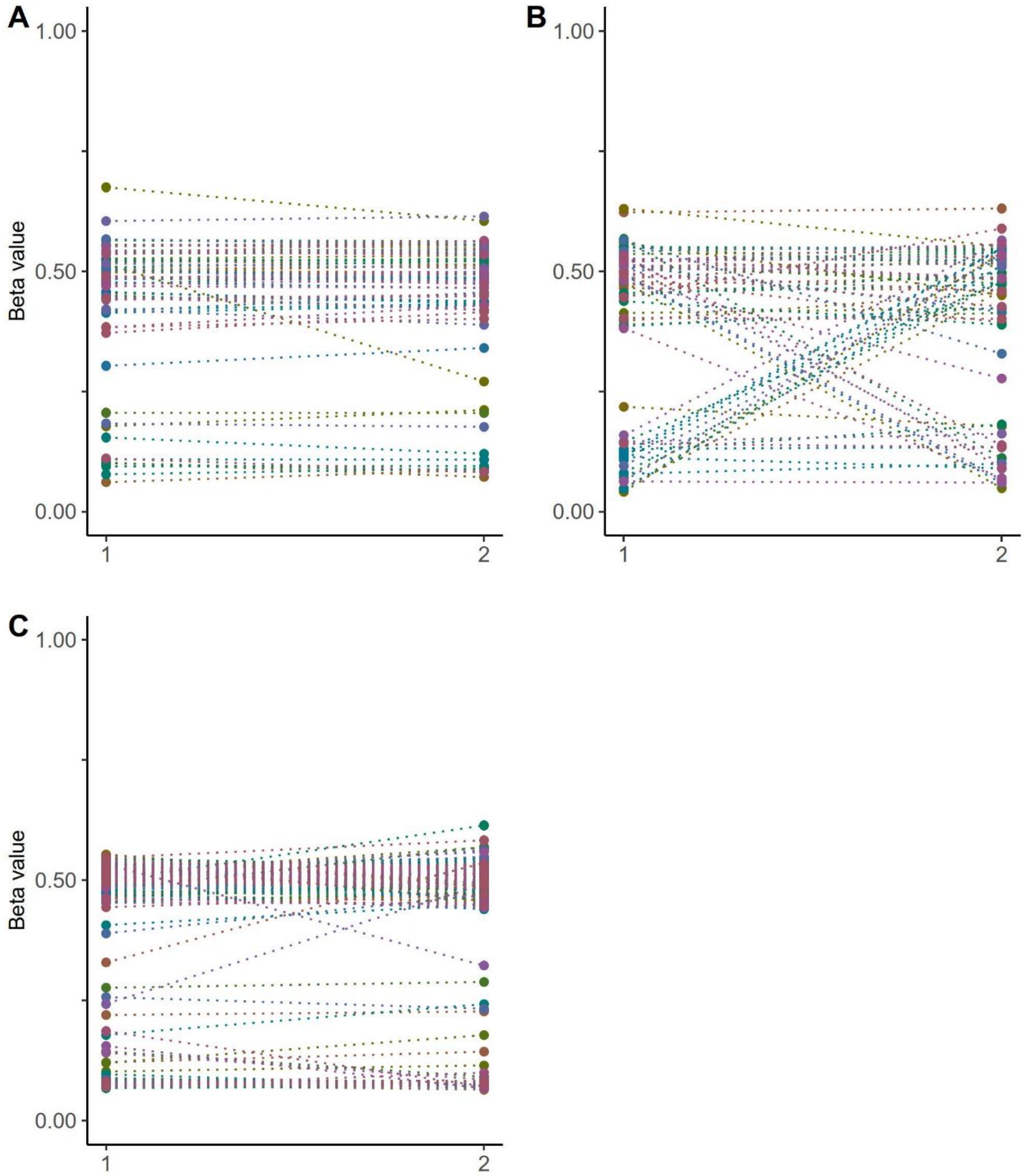


***Supplementary Figure 3.*** *Median methylation level of nc886 locus in A) monozygotic (66 pairs) and B) dizygotic twin pairs (66 pairs) from GSE100227 and C) monozygotic twin pairs from GSE61496 (156 pairs). In each figure one dot represents one individual, twin pairs have been connected with a dashed line.*

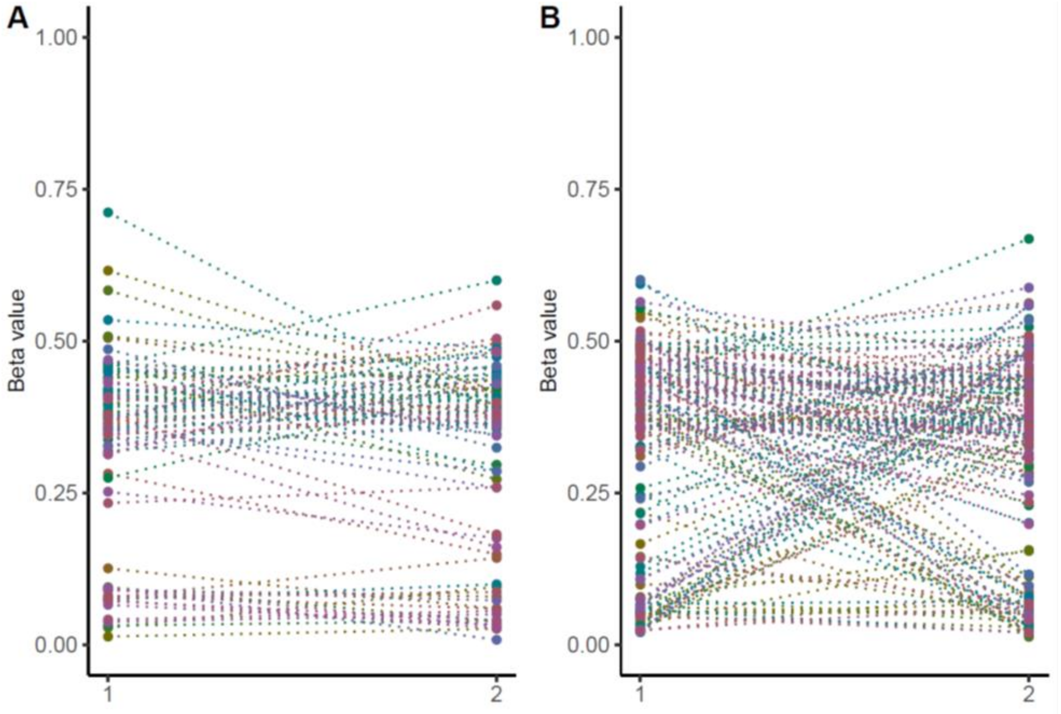


***Supplementary Figure 4.*** *Median methylation level of nc886 locus in SATSA in A) monozygotic twins (83 pairs) and B) dizygotic twins (156 pairs). In each figure one dot represents one individual, twin pairs have been connected with a dashed line.*

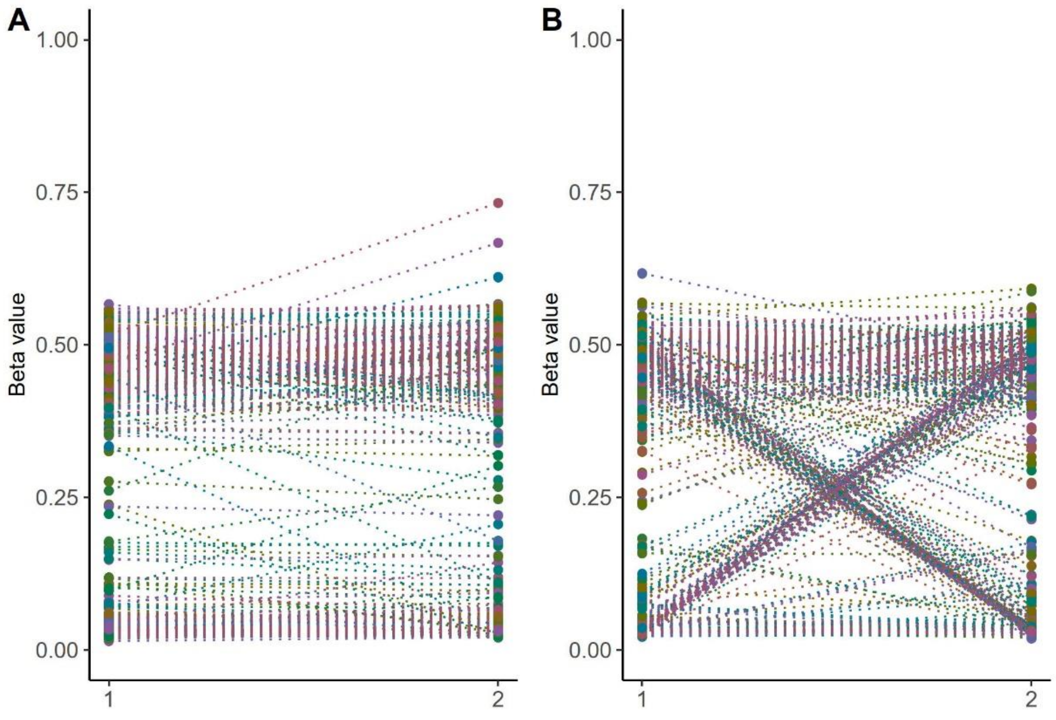


***Supplementary Figure 5****. Median methylation level of nc886 locus in FTC in A) monozygotic twins (519 pairs) and B) dizygotic twins (524 pairs). In each figure one dot represents one individual, twin pairs have been connected with a dashed line.*

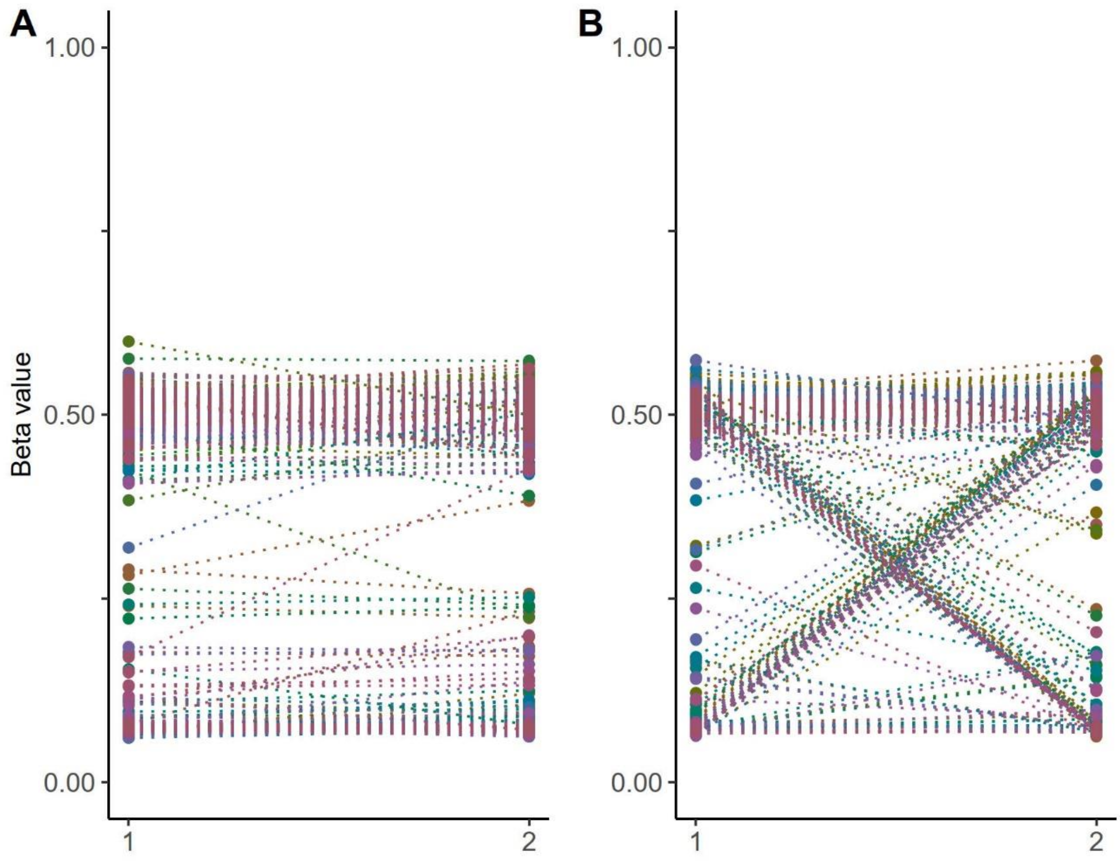


***Supplementary Figure 6.*** *Median methylation level of nc886 locus in A) monozygotic (426 pairs) and B) dizygotic (306 pairs) twin pairs (GSE 105018). In both figures each dot represents one individual, twin pairs have been connected with a dashed line.*

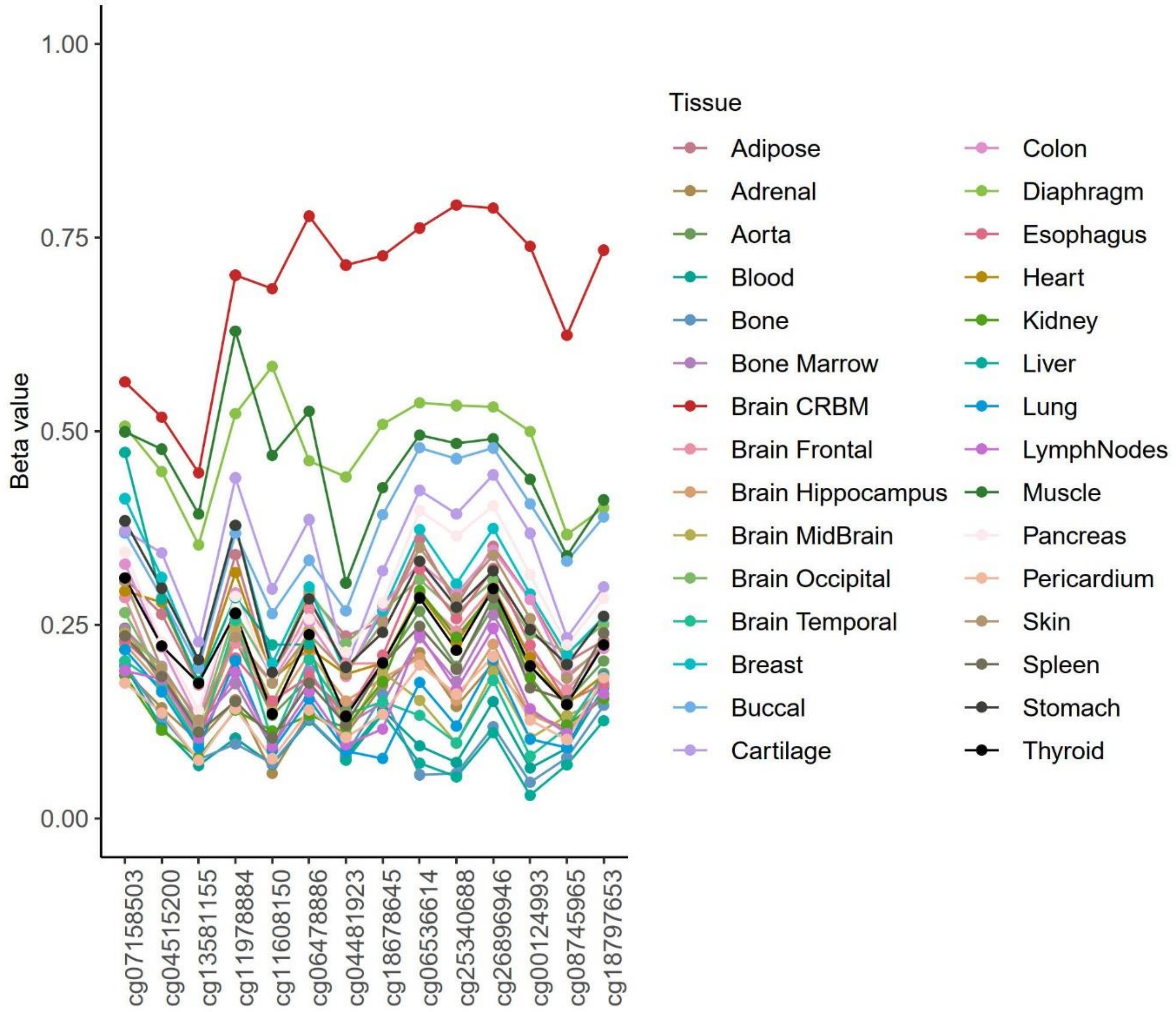


***Supplementary Figure 7.*** *Methylation level of the nc886 locus in 30 different tissues of a 112-year-old female (GSE64491). Tissues have been measured in duplicates or quadruplicates, in the figure each line is the median of replicates from the tissue.*

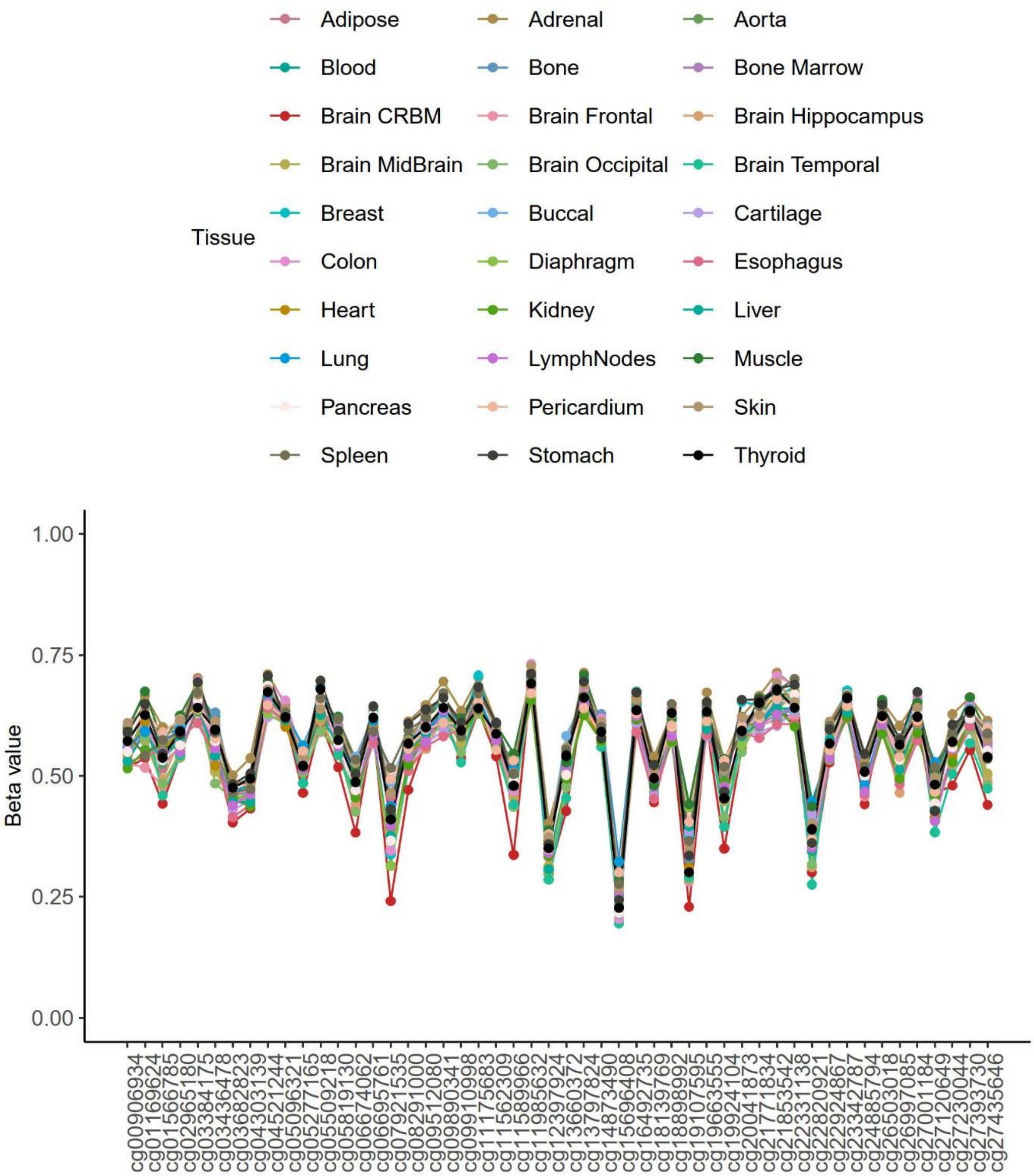


***Supplementary Figure 8.*** *Methylation level of PEG10 gene in 30 different tissues of a 112-year-old female (GSE64491). Tissues have been measured in duplicates or quadruplicates, in the figure each line is the median of replicates from the tissue. For five other known imprinted genes, DIRAS3, KCNQ10T1, MEG3, MEST, NAP1L5, and ZNF597, variation between tissues and between replicates was similar in magnitude as compared to PEG10 (Supplementary Table 5).*

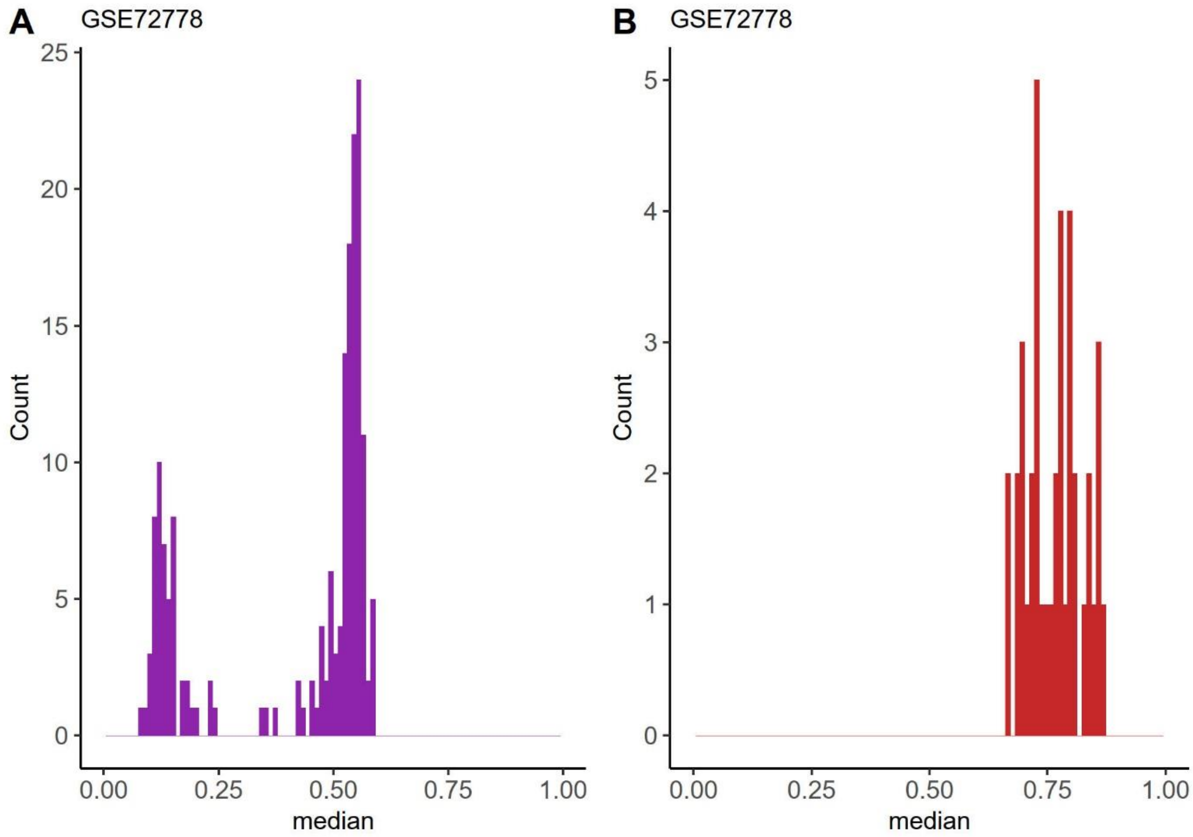


***Supplementary Figure 9.*** *Median methylation level of nc886 locus in A) five brain regions (frontal lobe, hippocampus, mid brain, occipital lobe and temporal lobe) and B) cerebellum (GSE72778).*

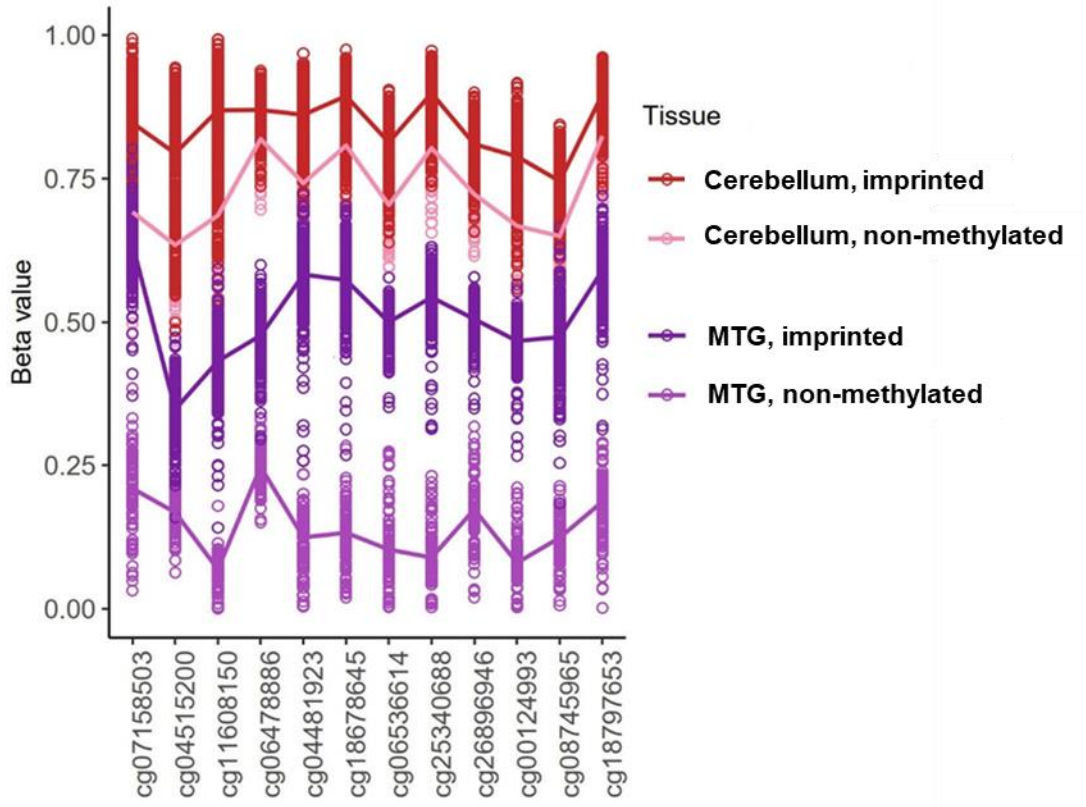


***Supplementary Figure 10**. Methylation level of the nc886 locus in cerebellum and middle temporal gyrus (MTG) (GSE134379). In the figure median of probes in each tissue is presented. Individuals were clustered based on methylation data from MTG as non-methylated (Non) and hemi-methylated (Hemi). Difference in methylation level in the cerebellum between non- and hemi-methylated individuals was statistically significant (Mann-Whitney U-test p-value < 0.001).*

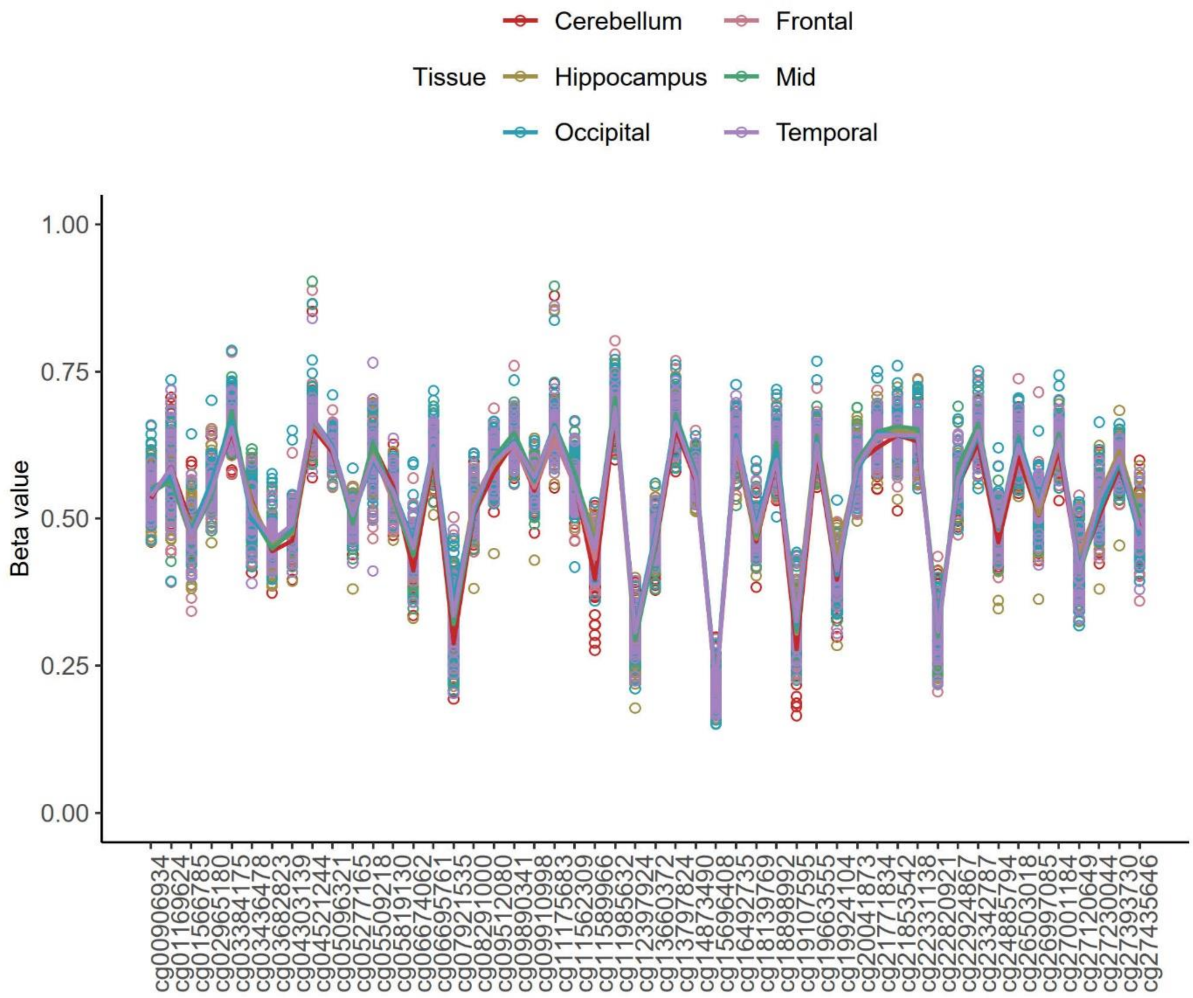


***Supplementary Figure 11.*** *Methylation level of PEG10 in cerebellum, frontal lobe, hippocampus, mid brain, occipital lobe and temporal lobe (GSE72778). Variation between brain regions was equally negligible also for DIRAS3, KCNQ10T1, MEG3, MEST, NAP1L5, and ZNF597 (data not shown).*

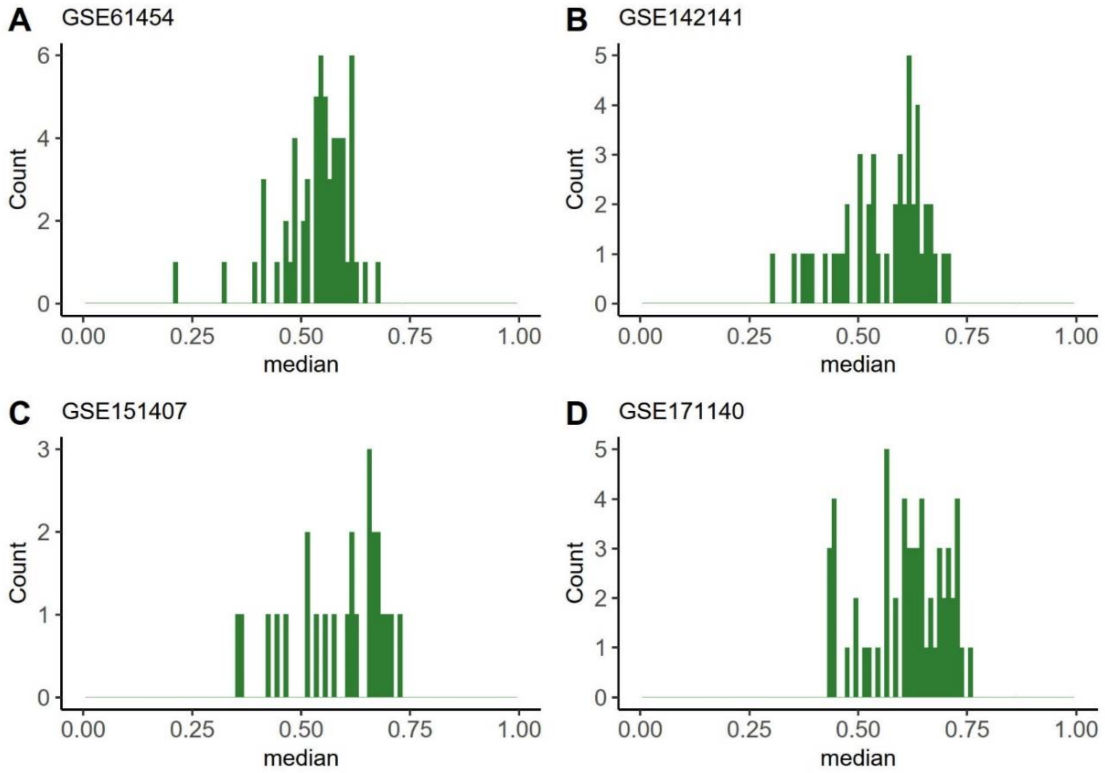


***Supplementary Figure 12***. *Methylation level at nc886 locus in muscle in four datasets A) GSE61454 n=60 (also presented in Figure 2), B) GSE142141 n=47, C)GSE151407 n=25, and D) GSE171140 n=57. Datasets GSE151407 and GSE171140 include partially same individuals, from these figures duplicates have been removed and each individual is presented only once. These two datasets also consist of baseline samples and samples taken after training intervention, only baseline samples have been included in these figures.*

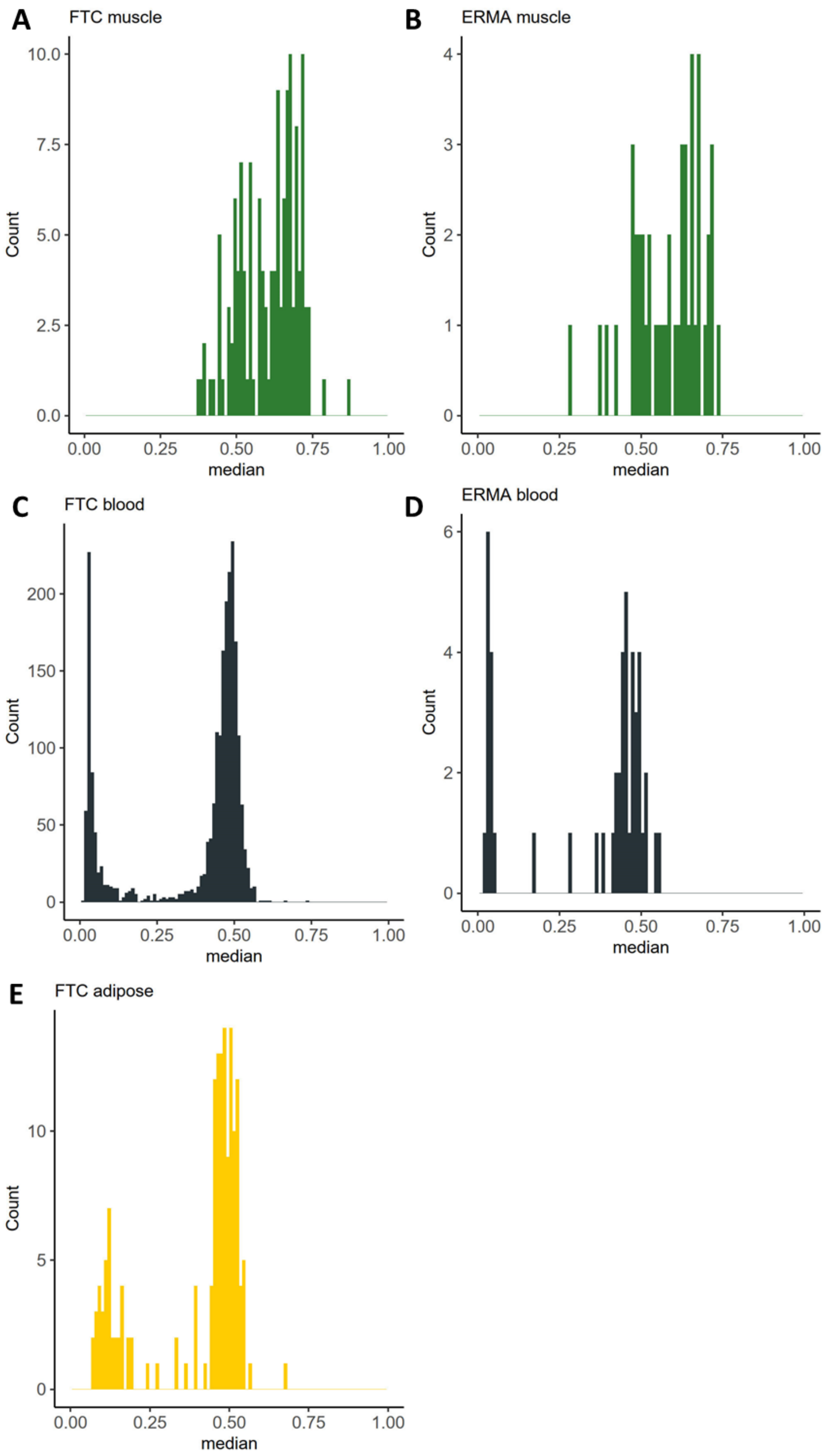


***Supplementary Figure 13.*** *Methylation level at nc886 locus in A) muscle in FTC, n=139, B) muscle in ERMA, n=47, C) blood in FTC, n=2240, D) blood in ERMA, n=47 and E) adipose tissue in FTC, n=160.*

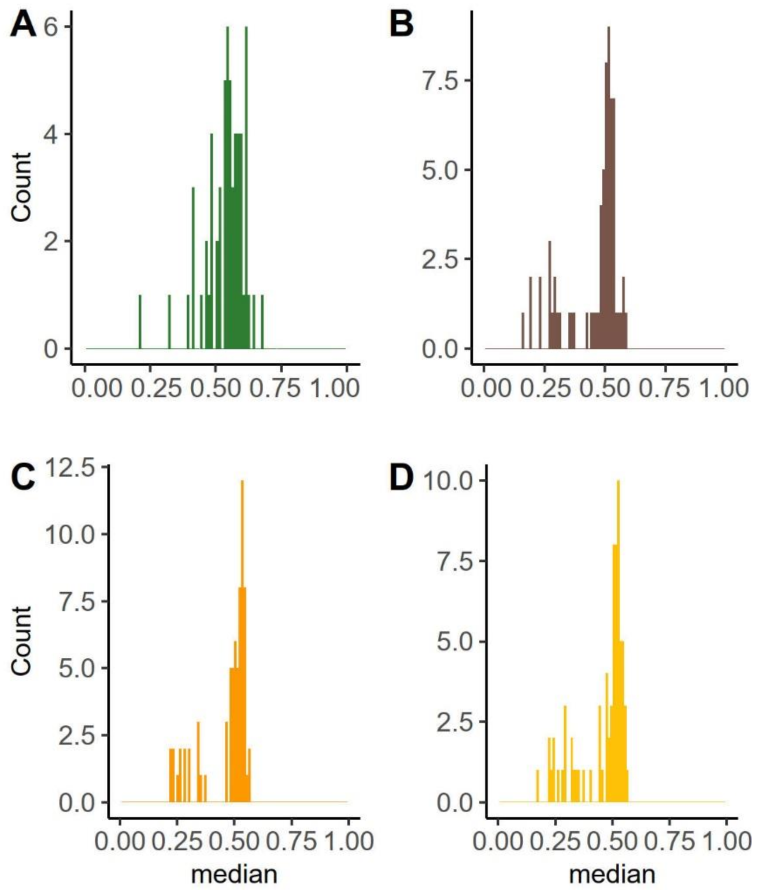


***Supplementary Figure 14***. *Methylation level at nc886 locus in dataset GSE61454 in A) muscle n=60 (also presented in Figure 2 and Supplementary Figure 12), B) liver n=67, C) visceral adipose tissue n=71, and D) subcutaneous adipose tissue n=71.*

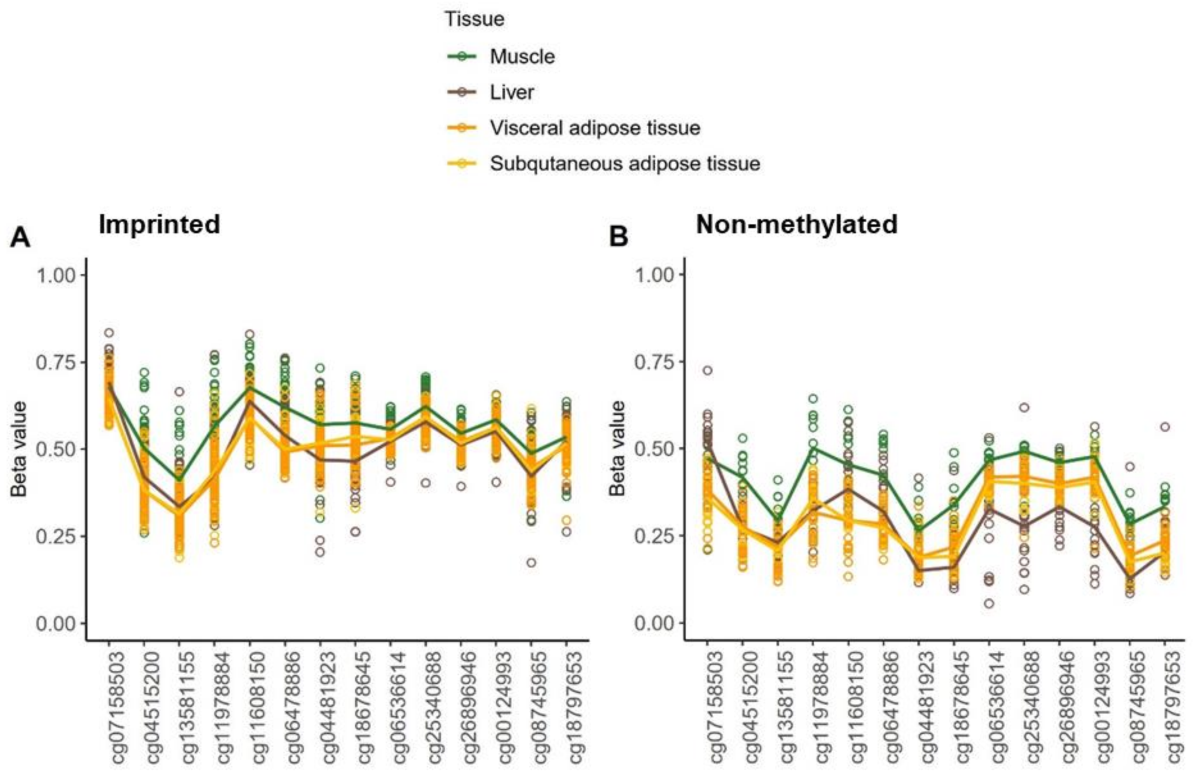


***Supplementary Figure 15**. Methylation level at nc886 locus in dataset GSE61454 in A) hemi-methylated individuals (as clustered based on liver and fat tissue) and B) non-methylated individuals. Despite the unimodal methylation level in muscle, we observed a difference in the methylation level at nc886 locus in muscle between non- and hemi-methylated individuals (Mann-Whitney U-test p-value < 0.001). Hemimethylated and nonmethylated in separate graphs to improve clarity.*

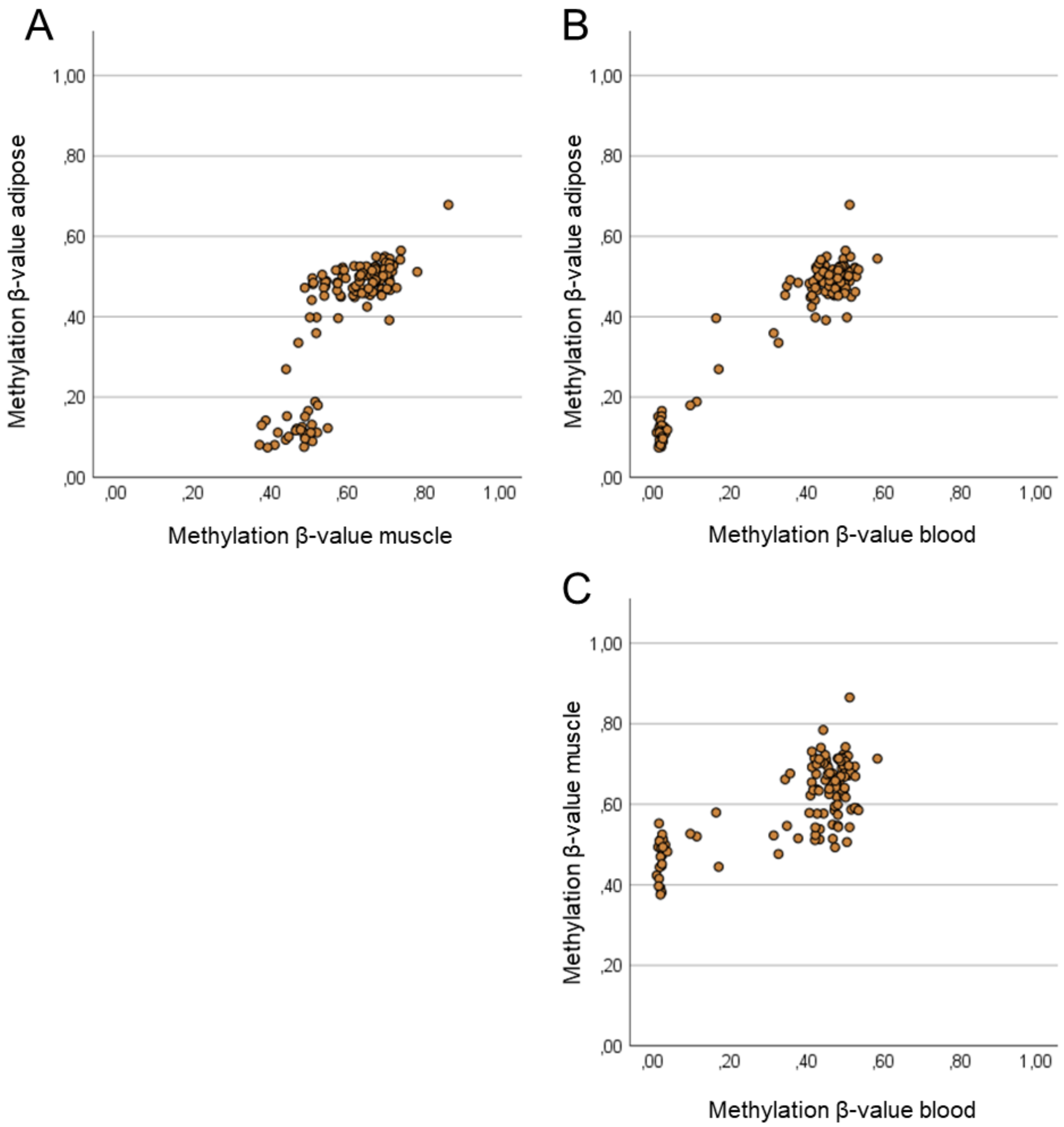


***Supplementary Figure 16.*** *Correlation of nc886 methylation levels in FTC (n=126) in A) between adipose tissue and muscle (ρ=0.719, p-value= 2.68*10$^{-21}$), B) adipose tissue and blood (ρ=0.660, p-value=4.26*10$^{-17}$) and C) muscle and blood (ρ=0.579, p-value=1.22*10$^{-12}$).*

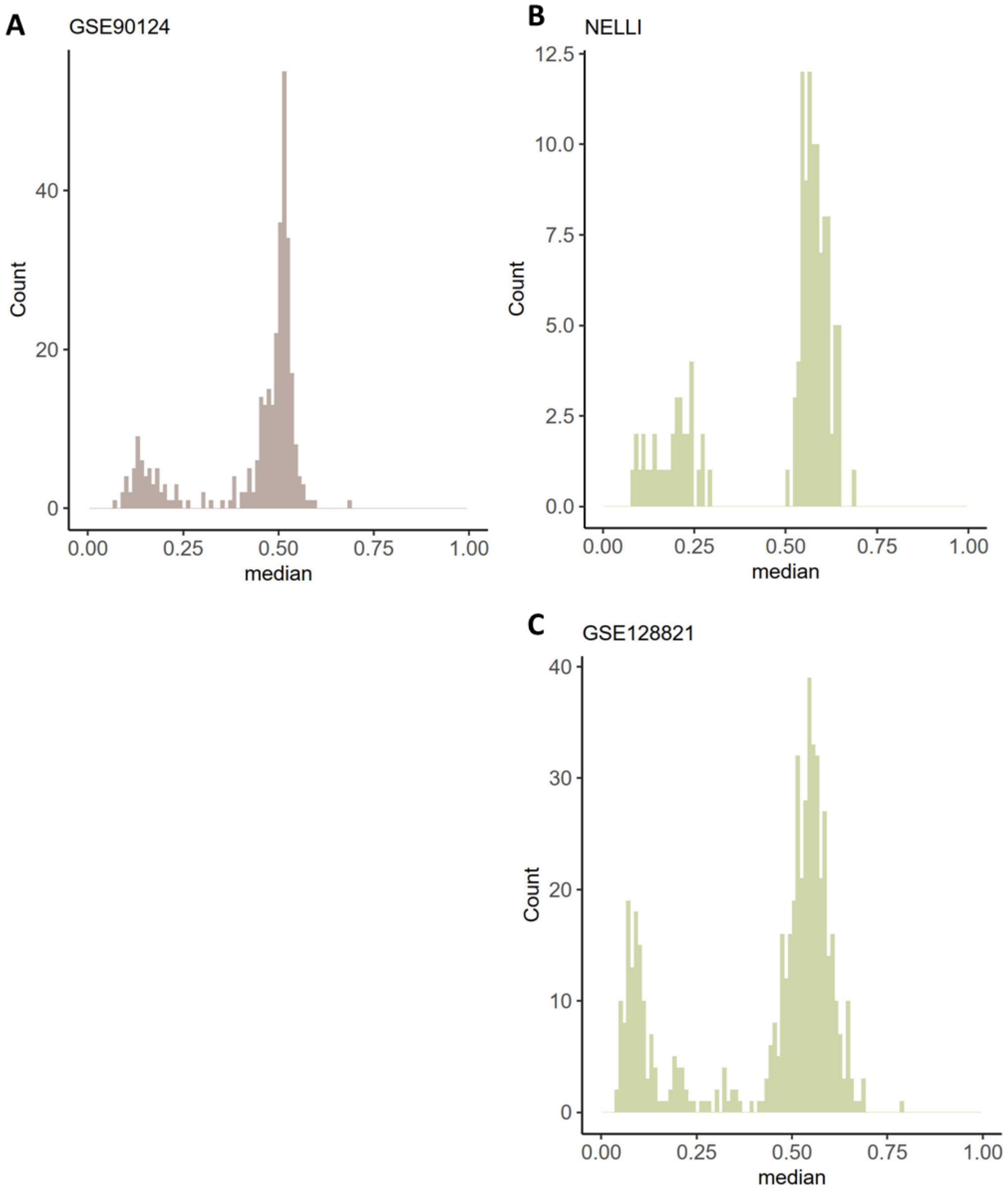


***Supplementary figure 17.*** *Methylation level at nc886 locus in A) skin (GSE90124), n=322 B) buccal swabs (NELLI), n=131 and C)buccal swabs (GSE128821), n=536.*

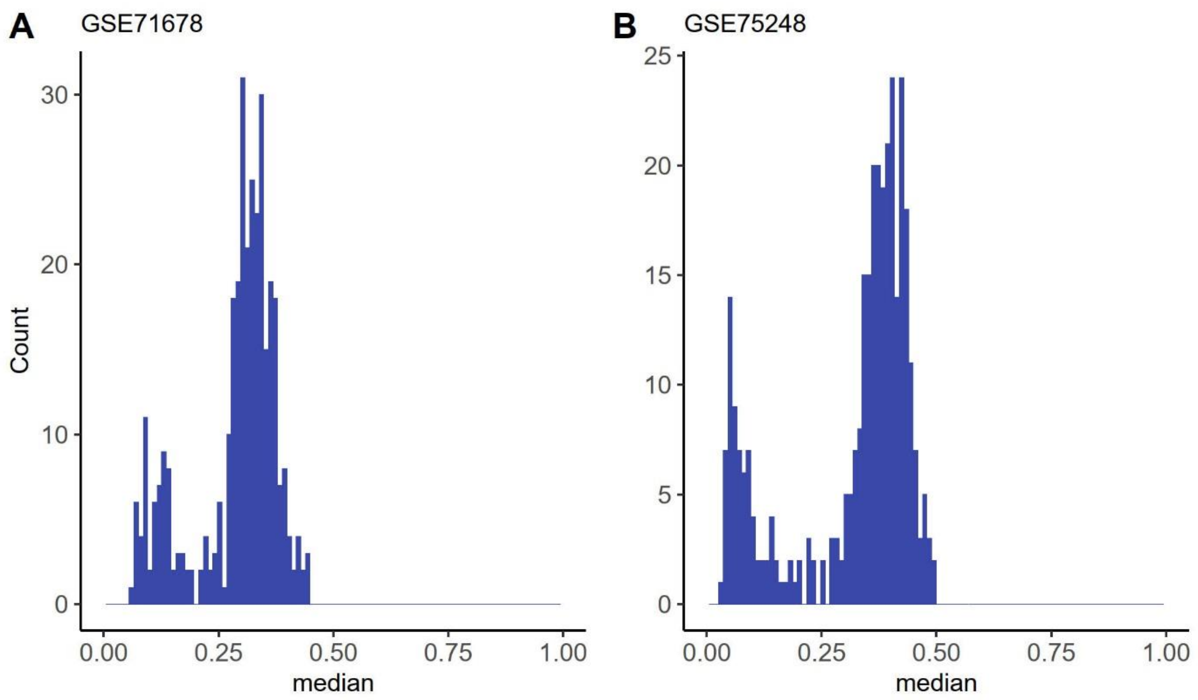


***Supplementary Figure 18***. *Methylation level of nc886 locus in placenta A) GSE71678 n= 343 and B) GSE75248 n= 335.*

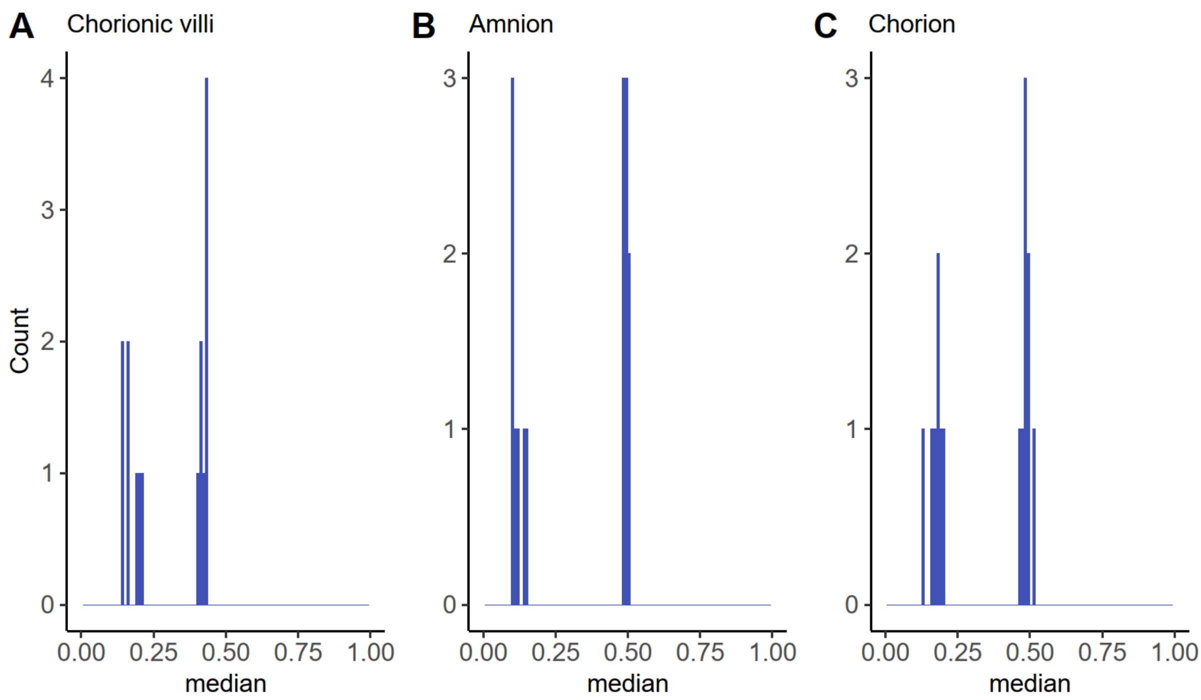


***Supplementary figure 19.*** *Methylation level of nc886 locus in dataset GSE115508 in A) placenta (chorionic villi) n=48 and corresponding B) amnion, n=15 and C) chorion, n=16.*

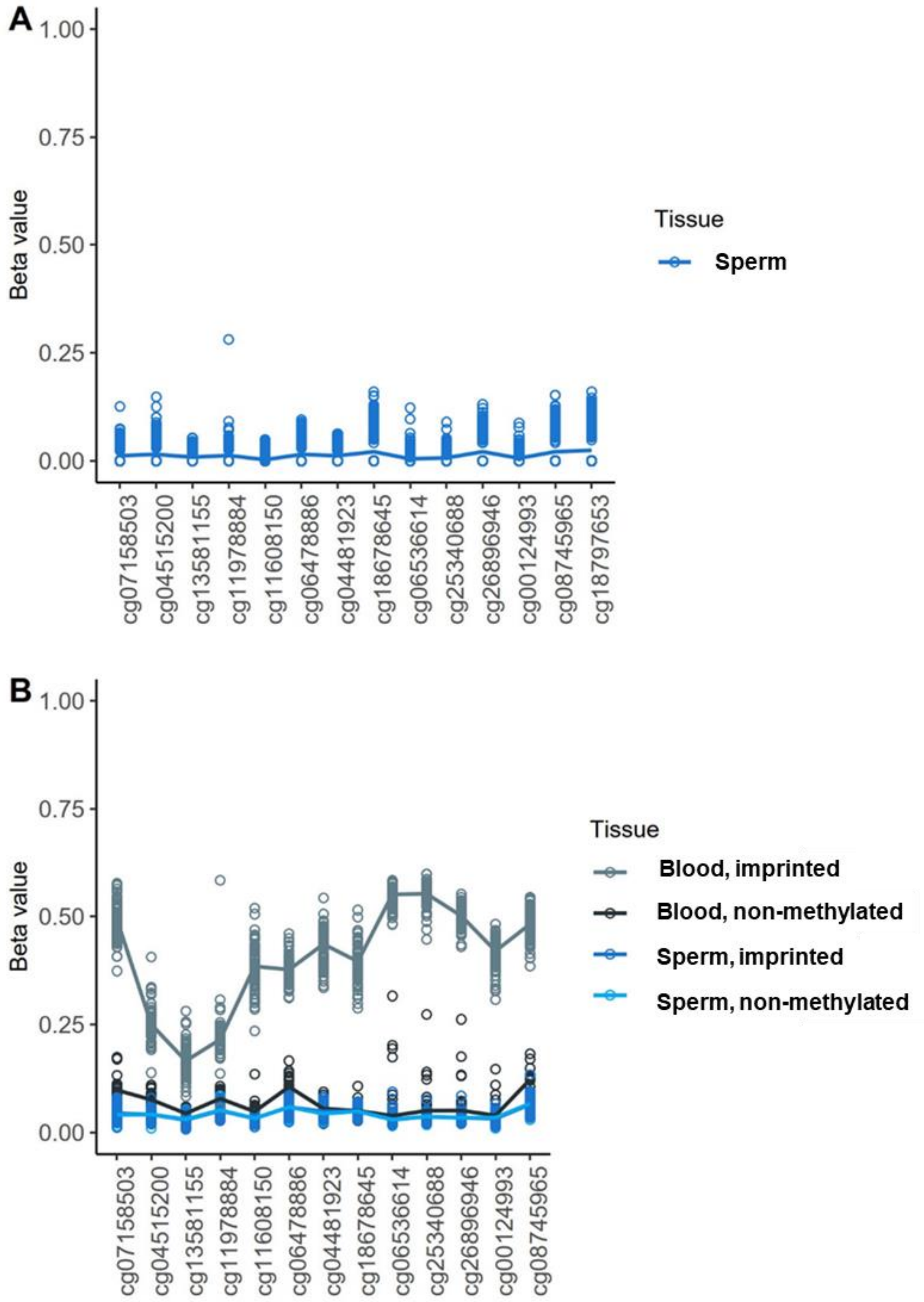


***Supplementary Figure 20.*** *Methylation level of nc886 locus in A) sperm, GSE114753, n=154 and B) blood and sperm, GSE149318, for sperm n=89, for blood n=90. For dataset GSE149318, we performed a clustering analysis to group individuals into imprinted and non-methylated according to blood nc886 methylation status. We observed no difference in sperm nc886 methylation level between imprinted and non-methylated individuals (Mann-Whitney U-test p-value > 0.05).*

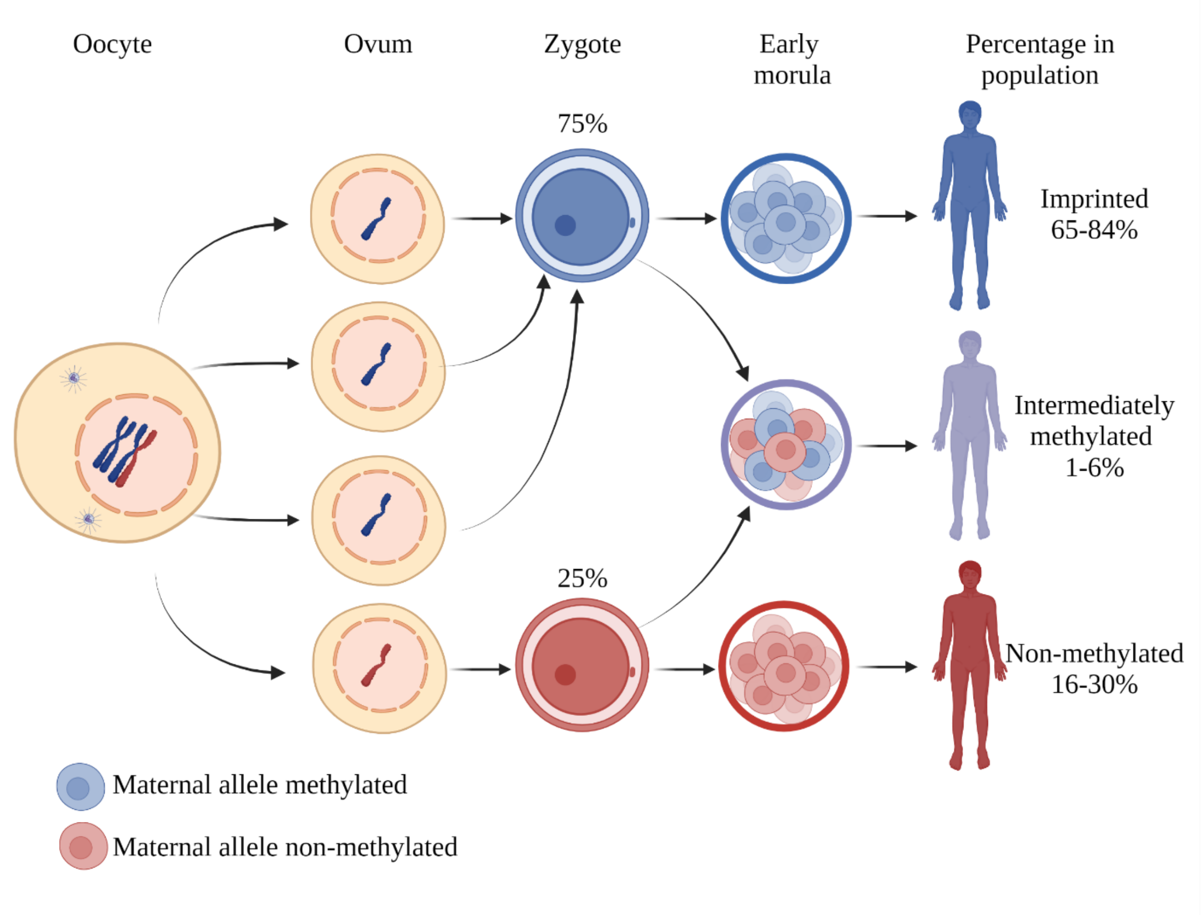


***Supplementary Figure 21.*** *Hypothesis on the establishment of nc886 imprinting. We hypothesize that the nc886 methylation status is established in the growing oocyte, similarly to other maternally imprinted genes. With a distribution of ¼ and ¾ in a biological system, one would expect there to be four entities present, when the distribution is established. In the case of establishing the imprint in the nc886 locus, by our hypothesis it should occur when there are four sister chromatids present in the developing oocyte. If by an unknown mechanism one of the chromatids is left unmethylated (and thus presumably permissive for gene expression), after completion of the meiosis and maturation of the ovum, each ovum would have a chance of 25% to harbour a non-methylated nc886 allele.*

*As demonstrated by data on MZ twin pairs, after fertilization, but prior to implantation, during the de- and re-methylation of embryonic genome, some individuals fail to maintain the established nc886 methylation status in all of their cells, leading to stochastic proportions of non-methylated and imprinted cells and thus intermediate nc886 methylation levels in tissue and individual level.*

*We hypothesize that the slight variation in the proportions of different nc886 methylation status groups could be explained by a survival advantage of one of the status groups during specific pregnancy conditions. We also cannot rule out that genetics could affect the maintenance of this imprinted during the global re- and demethylation of the embryonic genome, as we can see associations between the proportions of different status groups and ethnicity.*

*Paternal allele of nc886 is always non-methylated, as shown by the nc886 methylation level of close to 0 in sperm samples.*
